# Leptin antagonism improves Rett syndrome phenotype in symptomatic *Mecp2-*deficient mice

**DOI:** 10.1101/2023.02.03.526251

**Authors:** Yasmine Belaïdouni, Diabe Diabira, Pascal Salin, Mélanie Brosset-Heckel, Victoria Valsamides, Jean-Charles Graziano, Catarina Santos, Clément Menuet, Gary A. Wayman, Jean-Luc Gaiarsa

## Abstract

Rett syndrome (RTT) is a severe X-linked neurodevelopmental disorder caused by mutations in *MECP2*. Elevated circulating levels of the adipocyte hormone leptin are consistently observed in patients and in mouse models, yet their contribution to disease progression has remained unclear. Here, we show that reducing leptin signaling—either pharmacologically or genetically— significantly alleviates RTT-like phenotypes in *Mecp2*-deficient mice. In males, these interventions preserved general health, prevented weight loss, and improved breathing and locomotor functions. At the neuronal level, they restored excitatory/inhibitory balance in the hippocampus and somatosensory cortex and rescued hippocampal synaptic plasticity. In females, delaying the pathological rise of leptin levels postponed symptom progression. These findings uncover leptin as a key contributor to RTT pathophysiology and position leptin-targeted interventions as a promising therapeutic strategy for this currently untreatable disorder.

## INTRODUCTION

Rett syndrome (RTT; OMIM identifier #312750) is a rare non-inherited genetic neurodevelopmental disorder resulting from *de novo* mutations in the X-linked gene *MECP2* (methyl-CpG-binding protein 2) ^1^. The MECP2 protein binds to methylated DNA regulating the expression of thousands of genes and plays a crucial role in brain development and functioning ^2^. Affected individuals experience an apparently normal development until around 6–18 months of age. They subsequently undergo a rapid regression phase marked by the loss of purposeful hand movements and acquired speech, breathing irregularities, seizures and severe cognitive deficits ^3^. Several clinical features observed in humans with RTT are recapitulated in *Mecp2*-deficient mice, making these mouse models an essential tool for unraveling the cellular mechanisms underlying the disease and evaluating potential therapeutic interventions ^4^. Notably, studies have demonstrated that phenotypic rescue is achievable in *Mecp2-*deficient mice when the *Mecp2* gene is re-expressed even at late symptomatic stages ^5^. This finding indicates that RTT is not an irreversible condition, and it provides hope for therapeutic interventions.

Beyond *MECP2*, the expression of various molecular targets, including neurotrophic factors and neuromodulators, is dysregulated, some of which hold therapeutic potentials ^6^. Among these deregulated factors, elevated circulating levels of leptin have been observed in RTT patients and rodent models of the disease ^7–10^. Leptin, the product of the obese (*ob*) gene, is a circulating hormone secreted mainly from the white adipocytes. It crosses the blood brain barrier to act on the hypothalamus, where its main function is to regulate food intake ^11^. However, it is now clear that the actions of leptin extend beyond the hypothalamus, encompassing a wide range of biological roles. This pleiotropic hormone modulates motivation ^12^, cognitive functions ^13^, anxiety ^14^, the excitability of neuronal networks ^15^, epileptiform activities ^16^, breathing activity ^17^, locomotor activity ^18^ and more ^19,20^. Leptin also acts as an important neurotrophic factor during perinatal periods, promoting neuritic growth and synaptogenesis in various brain structures, including the hypothalamus ^21^ and hippocampus ^22–26^. Given these important physiological and developmental functions, dysregulation of leptin availability or signaling, either excess or deficiency, can have far reaching effects, including social and cognitive impairments, altered breathing activity, seizures susceptibility, locomotion and abnormal brain development ^16–18,24,27^. Consequently, the elevated levels of leptin observed in RTT patients ^7,8^ may contribute to the various disorders associated with Rett syndrome.

To address this hypothesis, we first confirmed the presence of elevated serum leptin levels in male and female *Mecp^2tm1–1Bird^* mice, hereafter referred to as *Mecp2*^-/y^ and *Mecp2*^+/-^ mice respectively. Subsequently, we evaluated the potential therapeutic effects of pharmacological antagonism of leptin and genetic reduction of leptin production in mitigating the deficits typically exhibited by this RTT mouse model. Our findings demonstrate that the pharmacological and genetic anti-leptin strategies prevent the degradation of health status, weight loss, and the progression of breathing and locomotor deficits in male *Mecp2*^-/y^ mice. At the neuronal level, these interventions rescue the excitatory/inhibitory balance in the hippocampus and somatosensory cortex and mitigate the impairment of synaptic plasticity in the hippocampus. Moreover, genetic reduction of leptin production delays the appearance of breathing and motor difficulties, and rescue the excitatory/inhibitory balance in the hippocampus of female *Mecp2*^+/-^ mice. These results indicate a role of leptin in the pathogenesis of RTT, providing valuable insights into potential therapeutic strategy for treating this syndrome.

## RESULTS

### Serum leptin levels and leptin mRNA expression are altered in male *Mecp2*^-/y^ mice

As a prerequisite for investigating the role of leptin in RTT, we assessed the circulating levels of leptin using ELISA, and the *Lep* mRNA expression levels from two main sources of leptin, the white adipose tissues (WAT) and gastrocnemius muscle, using qRT-PCR in male *Mecp2*^-/y^ and WT littermates. *Mecp2*^-/y^ mice showed elevated circulating leptin levels compared to their WT littermates. This difference reached statistical significance at postnatal day (P) 30 and 50 (**Figure 1A**). Despite weighting less than their WT littermates (**Figure 1B**), inguinal (**Figure 1C**), gonadic (1.03±0.06 vs 1.89±0.26 g/100g body weight, t=2.61, df=18, p=0.017, Unpaired *t*-test) and sub-cutaneous (0.46±0.07 vs 1.42±0.2 g/100g body weight, t=3.83, df=16, p=0.001, Unpaired *t*-test) WAT weighted significantly more in P50 *Mecp2*^-/y^ mice. The weight of visceral WAT was similar between WT and *Mecp2*^-/y^ littermates (**Figure 1C**). *Mecp2*^-/y^ mice also exhibited higher expression of leptin mRNA in gastrocnemius muscle (**Figure 1D**), as well as in visceral and inguinal WAT (**Figures 1E and F**). Taken together, these findings demonstrate that circulating leptin levels and leptin mRNA expression are elevated in male *Mecp2*^-/y^ mice when compared to their WT littermates, providing a foundation to investigate the role of leptin in RTT-associated symptoms.

**Figure 1:**
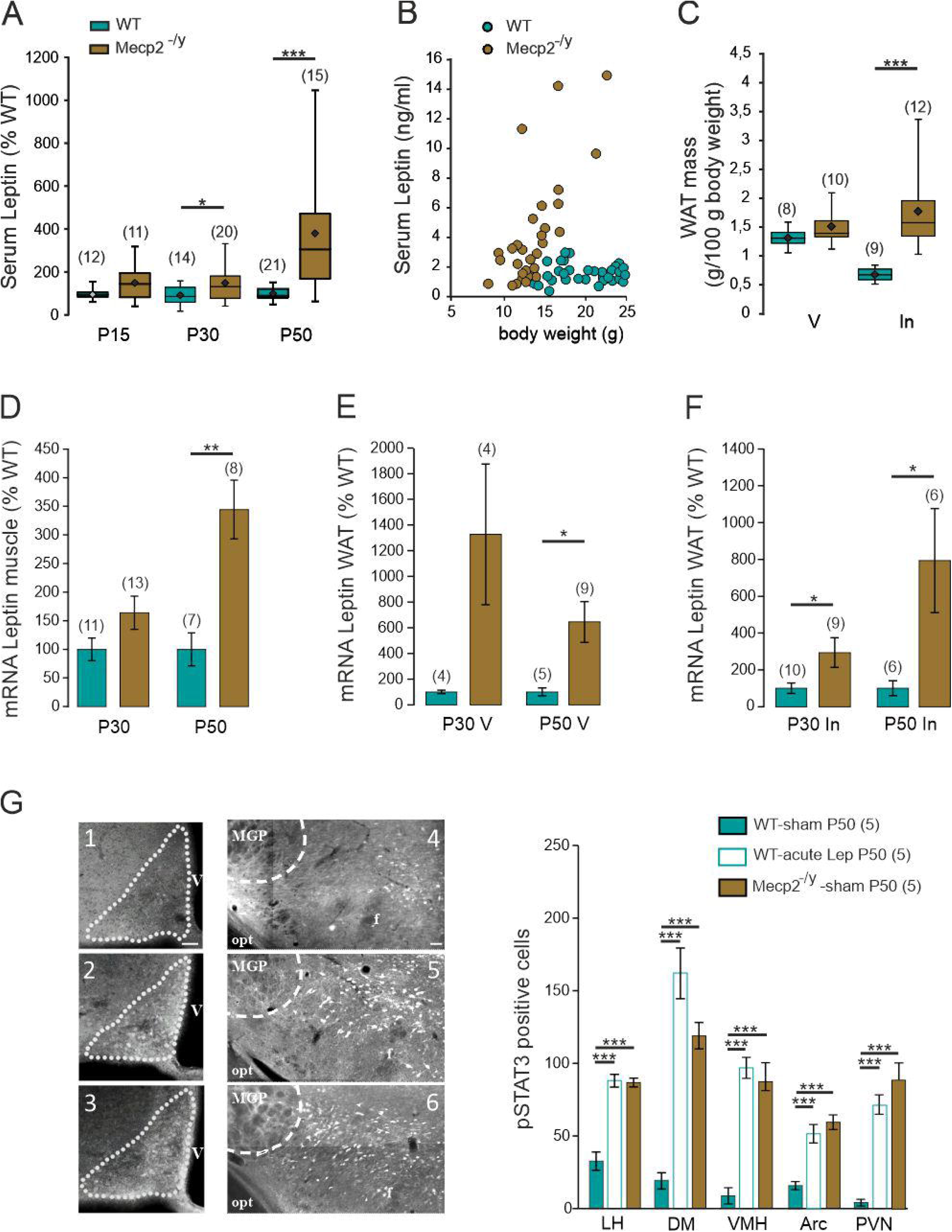
Circulating leptin levels, leptin mRNA expression and p-STAT3 immunostaining are elevated in symptomatic *Mecp2*^-/y^ mice. **A**) Box plots of serum leptin levels determined in blood samples taken at postnatal day (P) 15, 30 and 50, using ELISA kit and expressed as percentage of age matched wild type (WT) mice. **B)** Scatter plots of serum leptin levels as a function of mice body weight of P30-P50 WT and *Mecp2*^- /y^ mice. **C**) Box plots of normalized visceral (V) and inguinal (In) white adipose tissue (WAT) mass of P50 WT and *Mecp2*^-/y^ littermates. **D-F**) Mean±SEM plots of normalized leptin mRNA expression in gastrocnemius muscle (**D**), visceral WAT (**E**) and inguinal WAT (**F**) of P30 and P50 WT and *Mecp2*^-/y^ littermates. **(G)** G Representative microphotographs and quantification (mean±SEM) of pSTAT3 immunoreactivity in the delineated arcuate hypothalamic nucleus (bregma level -1.70) of WT Sham (1), WT Leptin (2) and KO Sham (3) and in the lateral hypothalamic area (bregma level -1.34) of WT Sham (4), WT Leptin (5) and KO Sham (6). V: third ventricle; Opt: optic tract; fr: fornix; delineated MGP: Medial globus Pallidus. Scale bars, 50 µmIn this and following figures, box plots represent quartiles, whiskers show data range, lozenges represent arithmetic means. Numbers in parentheses indicate the number of mice used. *P < 0.05, **P < 0.01, ***P < 0.001, two-tailed unpaired student *t*-test (C-F), One-way ANOVA followed by a Tukey’s multiple comparison (A, G).

We next aimed to determine whether hyperleptinemia results in sustained leptin receptor activation. To address this question, we used phosphorylated STAT3 (pSTAT3) as a surrogate marker of leptin receptor activation. Under basal conditions, *Mecp2*^-/y^ mice exhibited a greater number of pSTAT3-positive neurons across various hypothalamic nuclei compared to their WT littermates (**Figure 1G**), confirming previous findings ^28^. Interestingly, WT mice responded to acute sub-cutaneous injection of leptin (5µg/g) by increasing the number of pSTAT3 positive neurons in the hypothalamic nuclei to levels similar to those in *Mecp2^-/y^* mice (**Figure 1G**). Together, these results suggest that leptin signaling is enhanced in *Mecp2^-/y^*mice as a consequence of elevated circulating leptin levels, at least in hypothalamic nuclei.

### Leptin antagonism prevents the progression of breathing deficits in male *Mecp2*^-/y^ mice

Respiratory disorders have been well described in patients and rodent models of RTT ^29^. Leptin regulates breathing by acting on neurons of the brainstem respiratory network and carotid body cells ^17^. We therefore investigated the possible contribution of leptin to RTT-associated breathing disorders. We first characterized the respiratory activity of *Mecp2^-/y^* mice from our breeding colony. Consistent with previous studies ^30^, when compared with their age-matched WT littermates, P40 *Mecp2^-/y^* mice developed respiratory disorders characterized by increased number of apneas (**Figure 2A**), respiratory cycle duration variability expressed as breathing irregularity score (**Figure 2D**) and minute ventilation (**Figure 2F**). These breathing distresses worsen with age in *Mecp2^-/y^* mice (**Figures 2C and E**), while the respiratory parameters remained constant in WT mice (**Figures 2B and E**). Next, to test the possible role of leptin in RTT-associated breathing disorders, P40 WT mice received daily sub-cutaneous injection of leptin recombinant (5µg/g) during 10 consecutive days. Sham mice received the same volume of vehicle. We used plethysmography to record breathing activity of each mouse at P40, before starting treatment, and at P50, the last day of treatment. Leptin treatment of WT mice led to a significant increase in the number of apneas (**Figure 2B**) and breathing irregularity score (**Figure 2E**), while in sham WT mice these parameters remained unchanged (**Figures 2B and E**). The mean value of minute ventilation was not affected by the anti-leptin and leptin-treatment (**Figure 2G**).

**Figure 2:**
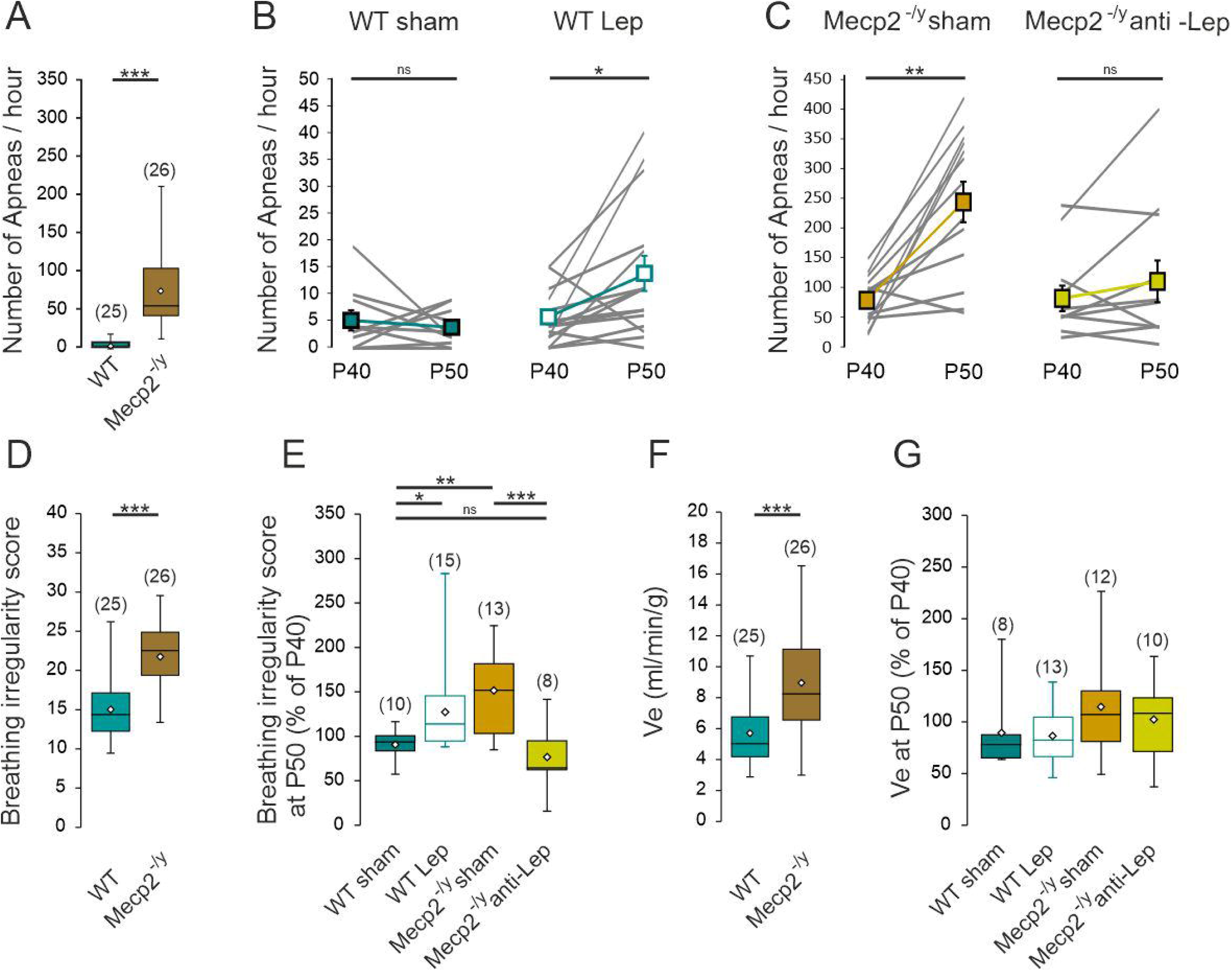
Leptin antagonism prevents the progression of breathing deficits of symptomatic *Mecp2*^-/y^ mice. **A)** Box plots of apnea frequency in P40 untreated WT and *Mecp2*^-/y^ mice. **B, C**) Plots of apnea frequency in treated WT (**B**) and *Mecp2*^-/y^ (**C**) mice. Lines indicate individual mice recorded the day before the treatment (P40) and at the end of the treatment (P50). Connected scares indicate mean±SEM. **D**) Box plots of breathing irregularity score in P40 untreated WT and *Mecp2*^-/y^ mice. **E)** Box plots of the percentage of change of breathing irregularity score in sham- and treated-WT and *Mecp2*^-/y^ mice. **F**) Box plots of minute ventilation in P40 untreated WT and *Mecp2*^-/y^ mice. **G**) Box plots of the percentage of change minute ventilation in sham- and treated-WT and *Mecp2*^-/y^ mice. Numbers in parenthesis indicate the number of mice used. \*\**P* < 0.01; \*\*\**P* < 0.001, Two-tailed paired t-test (B, C), Two-tailed unpaired t-test (A, D-G), One-way ANOVA followed by a Tukey’s multiple comparison (E, F).

We next investigated the effect of anti-leptin treatment (5µg/g, every other day during 10 days) on the breathing activity of the *Mecp2^-/y^* mice. While the sham and anti-leptin treated *Mecp2^-/y^* mice show similar phenotypic profile at the beginning of the treatment (**Figure 2C**), we found that the breathing distress worsen with age in sham treated *Mecp2^-/y^* mice, but remained constant in anti-leptin treated *Mecp2^-/y^* mice (**Figures 2C**). At the end of the treatment, P50 anti-leptin treated *Mecp2^-/y^* mice showed improved breathing parameters (apneas frequency and irregularity score) compared to sham *Mecp2^-/y^*mice (**Figures 2D & E**). The mean value of minute ventilation (**Figure 2F**) of P50 *Mecp2^-/y^*mice was not affected by the anti-leptin-treatment. We also tested the effect of chronic leptin treatment on *Mecp2^-/y^* mice and found no effect on their breathing parameters (**supplementary Figures S1A-C**). Altogether, these data suggest a contribution of leptin to the progression of breathing difficulties in symptomatic *Mecp2*^-/y^ mice.

### Leptin antagonism prevents the worsening of health condition of male *Mecp2^-/y^* mice

We next assessed the efficacy of the anti-leptin treatment on the overall health of the *Mecp2^-/y^* mice. The thoroughly described body weight loss of *Mecp2^-/y^*mice (**Figure 3A**) was abolished by the anti-leptin treatment (5µg/g, every other day during 10 days, from P40 to P50, **Figures 3B and C**). The leptin treatment (5µg/g during 10 consecutive days) decreased the body weight of the WT (**Figures 3C**) but not *Mecp2^-/y^* mice (**supplementary Figure S1D**). We also performed a scoring of the health condition of the mice. The parameters assessed include tremor, general aspect, spontaneous activity, limb grasp and posture. WT and *Mecp2^-/y^* mice showed significant difference at P40 (**Figure 3D**). *Mecp2^-/y^* mice showed altered general aspect, decreased spontaneous activity, and increased grasp and tremor (**Figure 3E**). The health scoring performed before starting treatment (P40) and on the last day of treatment (P50) revealed a deterioration in all parameters assessed in sham *Mecp2^-/y^* mice (**Figure 3G**). Worsening of posture, tremor and general aspect were prevented by the anti-leptin treatment while spontaneous activity and limb grasp showed no significant improvement (**Figure 3G**). Although it does not reach significance, this improvement in overall health after the anti-leptin treatment was accompanied by a decrease of early lethality, with 50% of the sham- and anti-leptin treated *Mecp2^-/y^*mice surviving after the postnatal day 48.5 and 59 respectively (**Figure 3F**). No lethality nor change in the health scoring were observed in WT mice treated with either vehicle or leptin (n=10 for both).

**Figure 3:**
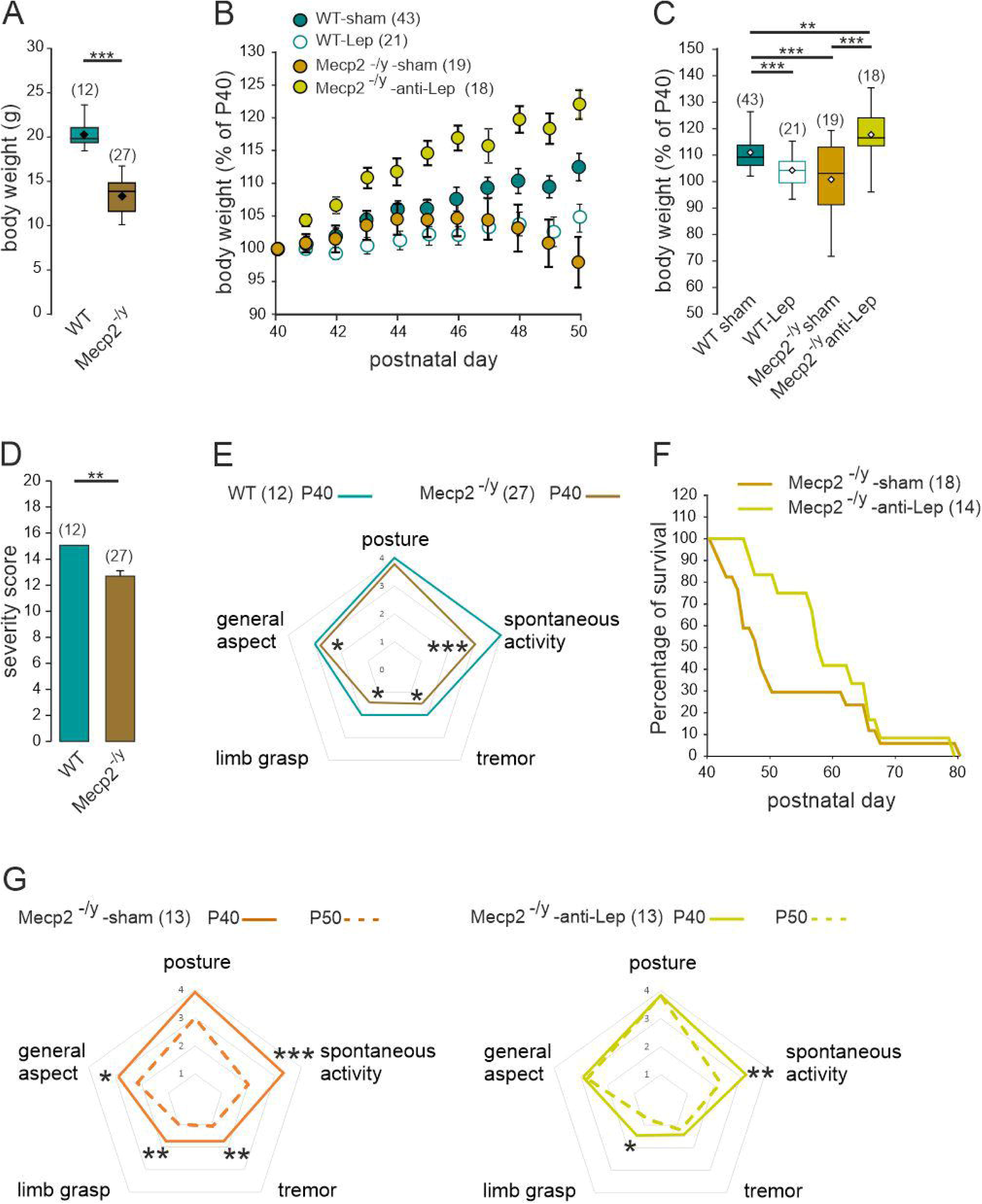
Leptin antagonism ameliorates the general health and lifespan of symptomatic *Mecp2^-/y^* mice. **A**) Box plots of the body weight of untreated P40 WT and *Mecp2^-/y^* mice. **B**) Body weight change (% of P40) as a function of age in sham and anti-leptin treated *Mecp2^-/y^*mice. The treatment started at P40 and ended at P50 (Mean±SEM). **C**) Box plot of the body weight change (% of P40) at P50 in sham and treated WT and *Mecp2*^-/y^ mice. **D**) Severity score of untreated P40 WT and *Mecp2^-/y^* mice. mean±SEM. **E**) Radar plot of the symptoms scored in P40 WT and *Mecp2^-/y^*mice. **F)** Plot of the percentage of surviving mice as a function of age. The treatment started at P40 and was maintained until the death of the animal. **G**) Radar plot of the symptoms scored in sham and anti-leptin treated *Mecp2*^-/y^ mice. Numbers in parenthesis indicate the number of mice used. \**P* < 0.05; \*\**P* < 0.01; \*\*\**P* < 0.001, Two-tailed unpaired t-test (A,C,D), One-way ANOVA followed by a Tukey’s multiple (C), Mann Whitney U test (E, G), Kaplan-Meier log rank test (F).

Although the underlying mechanisms require further clarification, leptin has been reported to modulate locomotor activity ^18^. We therefore assessed the effect of the anti-leptin treatment on the locomotor activity and motor coordination of *Mecp2^-/y^* mice in the open field and the accelerating rotarod tests. In the open field test, the distance traveled by the *Mecp2^-/y^* mice was reduced compared to their WT littermates (**Figures 4A**). This locomotor deficit worsened with age in *Mecp2^-/y^* mice (**Figures 4B**). The 10-day anti-leptin treatment (5µg/g, from P40 to P50) prevented the degradation of locomotor activity observed in the sham *Mecp2^-/y^* mice (**Figures 4B and 4C**). Moreover leptin-treated WT mice performed similarly to their sham littermates (**Figure 4C**). In the accelerating rotarod test, *Mecp2^-/y^* mice showed a significant decrease in the latency to fall compared to WT littermates (**Figure 4D**). This phenotype did not worsen with age (**Figures 4E**). In the accelerating rotarod test, neither the leptin-nor the anti-leptin treatment affected the performances of respectively WT and *Mecp2^-/y^*mice (**Figure 4F**). Overall, these observations show that the anti-leptin treatment ameliorates the general health condition, (i.e., posture, tremor, general aspect) and locomotor activity of the symptomatic *Mecp2^-/y^* mice.

**Figure 4:**
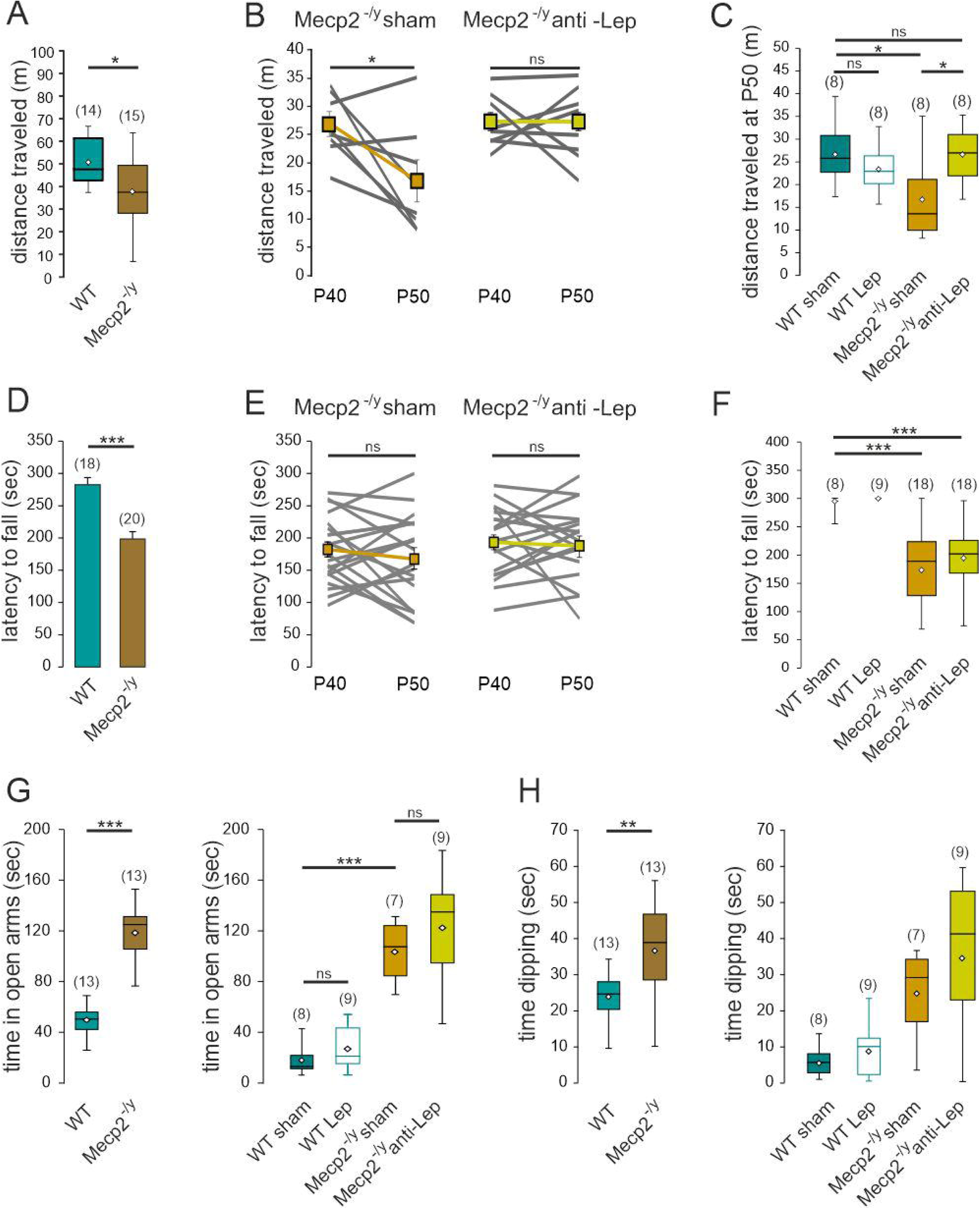
Leptin antagonism prevents the worsening of locomotor deficit but does not improve motor coordination nor anxiety alterations of symptomatic *Mecp2^-/y^* mice. A) Box plots of the distance traveled by untreated P40 WT and *Mecp2^-/y^* mice in an open field. **B**) Plots of distance traveled by sham and anti-leptin *Mecp2*^-/y^ mice in an open field. Connected points indicate individual mice recorded the day before the treatment (P40) and at the end of the treatment (P50). Connected squares indicate mean±SEM. **C**) Box plots of the distance traveled in an open field by P50 sham and treated WT and *Mecp2*^-/y^ mice at the indicated genotype and treatment. **D**) Mean ± SEM plots of the latency to fall of untreated P40 WT and *Mecp2^-/y^* mice in the accelerating rotarod. **E**) Plots of latency to fall by sham and anti-leptin *Mecp2*^-/y^ mice. Connected points indicate individual mice recorded the day before the treatment (P40) and at the end of the treatment (P50). Connected squares indicate mean±SEM. **F**) Box plots of the latency to fall by P50 sham and treated mice at the indicated genotype and treatment. **G, H**) Box plots of the time spent in open arms (**G**) and dipping (**H**) by untreated P40 WT and *Mecp2^-/y^* mice, and P50 WT- and *Mecp2^-/y^*-sham and treated mice in an elevated plus maze. Numbers in parenthesis indicate the number of mice used. \**P* < 0.05; \*\**P* < 0.01; \*\*\**P* < 0.001, two-tailed unpaired *t*-test when comparing untreated mice (A,C,F,G,H,), two-tailed paired *t*-test (B,E), Mann Whitney U test (D), Fisher exact test (J), Mann Whitney test (D), One-way ANOVA followed by a Tukey’s multiple comparisons when comparing treated mice (C,F,G,H,I).

### Leptin antagonism does not affect anxiety or seizures susceptibility in male *Mecp2*^-/y^ mice

Leptin has been reported to exert anti-depressant and pro-cognitive like effects in rodents ^13,14^. We therefore investigated the possible contribution of leptin to cognitive behavioral deficits repeatedly reported in RTT models. Our first set of behavioral tests examined anxiety, cognition and social interaction in P50 WT and *Mecp2^-/y^* mice from our breeding colony. We first compared the spontaneous exploration in the elevated plus maze (EPM) as a measure of anxiety and found that *Mecp2^-/y^* mice spent more time in the open arms than their WT littermate (**Figure 4G**). The mean dipping time was also increased in *Mecp2^-/y^* mice (**Figure 4H**) a behavior also consistent with decreased anxiety. However, no genotype effects were found in tests used to evaluate object recognition and social behavior (**supplementary Figure S2**). We therefore limited the study of the role of leptin to the EPM. Leptin-treated WT and anti-leptin treated-*Mecp2^-/y^* mice performed similarly to their vehicle-treated littermates in the EPM, i.e., they stayed for a similar amount of time in the open arms and exhibited similar mean dipping time (**Figures 4H and G**). These observations therefore show that the anti-leptin treatment does not modify the anxiety-related behavior of symptomatic *Mecp2^-/y^* mice.

### Leptin antagonism restores the E/I balance in CA3 hippocampal neurons and layer 5 somatosensory cortical neurons, and mitigates the hippocampal synaptic plasticity in male *Mecp2*^-/y^ mice

We then assessed the potential benefits of anti-leptin treatment at the neuronal and cellular levels. Alteration in the synaptic excitation/inhibition (E/I) balance is a widespread feature of neuronal networks in *Mecp2*-deficient mice ^31^. Leptin has been reported to modulate the development and functioning of both glutamatergic and GABAergic synapses ^13,23–26^, raising the possibility that elevated leptin levels might contribute to the altered E/I balance in *Mecp2*^-/y^ mice. We performed whole cell patch clamp recordings of CA3 pyramidal neurons on acute hippocampal slices obtained from P50 WT and *Mecp2*^-/y^ mice. We found that, as already reported ^32^, the E/I balance was increased in *Mecp2*^-/y^ compared to their WT littermates (**Figures 5A and B**). The 10-day leptin treatment (5µg/g, from P40 to P50) led to a significant increase in the E/I balance in P50 WT mice (**Figure 5B**), while having no effect in *Mecp2*^-/y^ mice (**supplementary Figure S1G**). Conversely, the 10-day anti-leptin treatment (5µg/g from P40 to P50) restored the E/I balance in *Mecp2*^-/y^ mice (**Figure 5B**). We also asked whether the anti-leptin treatment could correct altered E/I balance in cortical neurons of *Mecp2*^-/y^ mice. Whole cell patch clamp recordings of somatosensory layer 5 cortical neurons revealed that E/I balance was increased in *Mecp2*^-/y^ mice compared to their WT littermates (**Figure 5C**) and normalized by the 10-day anti-leptin treatment (5µg/g from P40 to P50) (**Figures 5C**). Altogether, these results show that leptin contributes to the E/I imbalance in symptomatic *Mecp2*^-/y^ mice.

**Figure 5:**
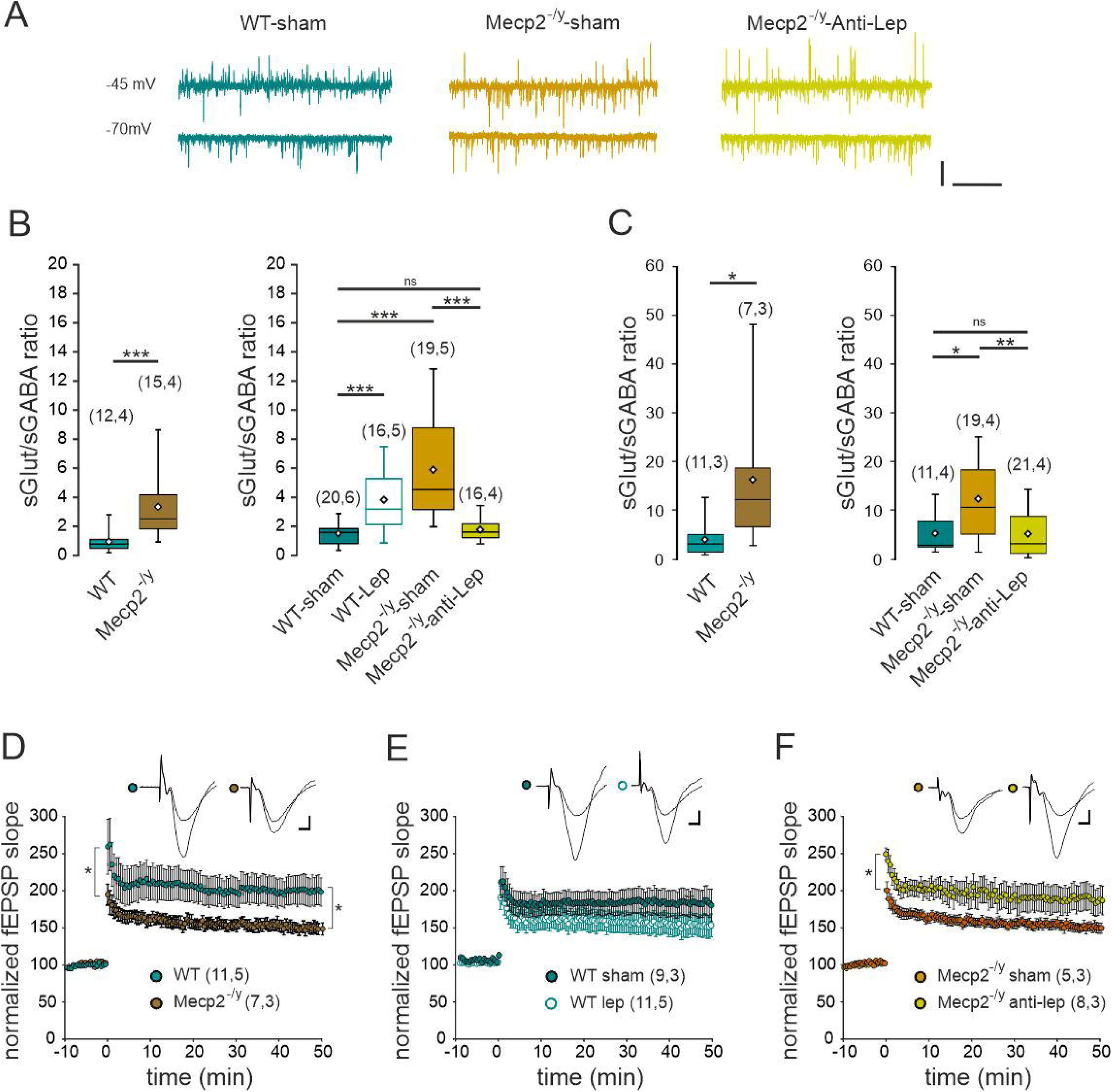
Leptin contributes to hippocampal and cortical synaptic alterations of symptomatic *Mecp2*^-/y^ mice. **A)** Examples of whole-cell recordings performed on CA3 pyramidal neurons at a holding potential of −45 and -70 mV at P50. The glutamatergic synaptic currents are inwards and the GABAergic synaptic currents are outwards. Scale bars: 100pA, 2 sec. **B, C)** Box plots of the ratio frequency of spontaneous glutamatergic and GABAergic postsynaptic currents recorded from P50 untreated and treated WT and *Mecp2*^-/y^ CA3 pyramidal neurons (**B**) and layer V somatosensory pyramidal neurons (**C**). **D-F**) LTP was recorded for 50 min following tetanic stimulation at the Schaffer collateral-CA1 synapses of acute hippocampal slices from symptomatic (P50) untreated (**D**) and treated (**E, F**) *Mecp2*^-/y^ and WT mice. Insets show examples of fEPSP. Scale bar: 100µV, 3ms. Numbers in parenthesis indicate the number of slices recorded and mice used. Numbers in parenthesis indicate the number of cells recorded and mice used. \*\**P* < 0.01; \*\*\**P* < 0.001, two-tailed unpaired *t*-test when comparing untreated mice (B,C), One-way ANOVA followed by a Tukey’s multiple comparison when comparing treated mice (B,C). Repeated measure ANOVA (D-F).

Impairment of synaptic plasticity is another common feature of mice RTT models ^33^ and leptin has been reported to modulate the strength of glutamatergic synapses ^27^. We therefore examined the effect of leptin and leptin antagonist treatment on long term potentiation (LTP) at the hippocampal Schaffer collaterals-CA1 synapses of P50 WT and *Mecp2*^-/y^ mice. As already reported ^33^, the magnitude of early (0-2 min post-tetanus) and late (45-50 min post-tetanus) LTP were reduced in *Mecp2*^-/y^ mice compared to WT littermates (**Figure 5D**). Leptin treatment of WT mice (5µg/g during 10 consecutive days from P40) attenuated the magnitude of late and early LTP compared to sham-treated mice but these differences did not reach significance (**Figure 5E**). Conversely, the anti-leptin treatment (5µg/g every other day during 10 days from P40) increased the magnitude of early and late LTP in *Mecp2*^-/y^ mice (**Figure 5H**). Altogether, these data demonstrate the major contribution of leptin elevation to the hippocampal LTP impairment in symptomatic *Mecp2*^-/y^ mice.

Studies have shown that the brain derived neurotrophic factor (BDNF) levels are reduced in *Mecp2*-deficient mice and that their recovery would be a potential candidate for the treatment of RTT ^34^. Leptin modulates BDNF expression in several brain structures including the hypothalamus ^35^ and hippocampus ^36^. To determine whether the anti-leptin treatment modulates BDNF expression, we quantified the levels of mBDNF and proBDNF proteins in P50 hippocampi using ELISA. As already reported ^34^, the level of mBDNF was significantly lower in *Mecp2*^-/y^ mice compared to their WT littermates (**supplementary Figure S3**). The level of proBDNF in *Mecp2*^-/y^ mice was not different from that of WT mice. The anti-leptin treatment (5µg/g every other day during 10 days from P40) did not restore the level of mBDNF in *Mecp2*^-/y^ mice (**supplementary Figure S3**). Therefore, BDNF is unlikely to mediate the effects on the ant-leptin treatment in the hippocampus of *Mecp2*^-/y^ mice.

### RTT-associated symptoms are improved in male and female leptin haplo insufficient *Mecp2*-deficient mice

The data presented above show that leptin antagonism effectively mitigates the deterioration of, or even rescues, certain symptoms observed in symptomatic *Mecp2*^-/y^ mice. However, prolonged administration of a leptin antagonist presents challenges, as it may potentially lead to metabolic and/or immune disturbances. To reduce this drawback and further evaluate the possible contribution of leptin in RTT phenotype, we generated *Mecp2*^-/y^ mice that are haplo insufficient for leptin with the aim of restoring the physiological levels of leptin in *Mecp2*-deficient mice. Leptin haplo insufficient *Mecp2*^-/y^ mice were generated by backcrossing *Mecp2^+/-^*female mice with heterozygous leptin deficient male mice (*ob^+/-^*). The resulting double transgenic *Mecp2*^- /y^*;ob^+/-^* mice showed normalized leptin levels at P40-50 (**Figure 6A**). We next examined whether *Mecp2*^-/y^*;ob^+/-^* mice exhibited any improvements at advanced symptomatic stage. Notably, the progressive deterioration in breathing activity typically observed in *Mecp2*^-/y^ mice was prevented in *Mecp2*^-/y^*;ob^+/-^* mice (**Figures 6B and C**). In addition, *Mecp2*^-/y^*;ob^+/-^* mice displayed significantly higher body weight compared to *Mecp2*^-/y^ littermates, with this difference becoming increasingly pronounced as the disease progressed (**Figures 6D**). Whole cell recordings performed on hippocampal slices revealed that the E/I balance (**Figures 6E**) was improved in CA3 hippocampal neurons of *Mecp2*^-/y^*;ob^+/-^* mice. Field recordings on hippocampal slices further revealed that the magnitude of early and late LTP at the Schaffer collateral-CA1 synapses was increased in *Mecp2*^-/y^*;ob^+/-^* mice (**Figure 6F**). Analysis of the health condition of the *Mecp2*^- /y^*;ob^+/-^* mice revealed some improvement at P40 but not at P50 (**Figure 6G**). At P40, *Mecp2*^- /y^*;ob^+/-^* mice showed fewer tremor and limb grasping compared to the *Mecp2*^-/y^ mice (**Figure 6H**). The life expectancy of *Mecp2*^-/y^*;ob^+/-^* mice was however not different compared to their *Mecp2^-/y^* littermates (**Figure 6I**). Overall, these data show that reducing leptin production improves some RTT-associated symptoms in *Mecp2^-/y^* mice.

**Figure 6:**
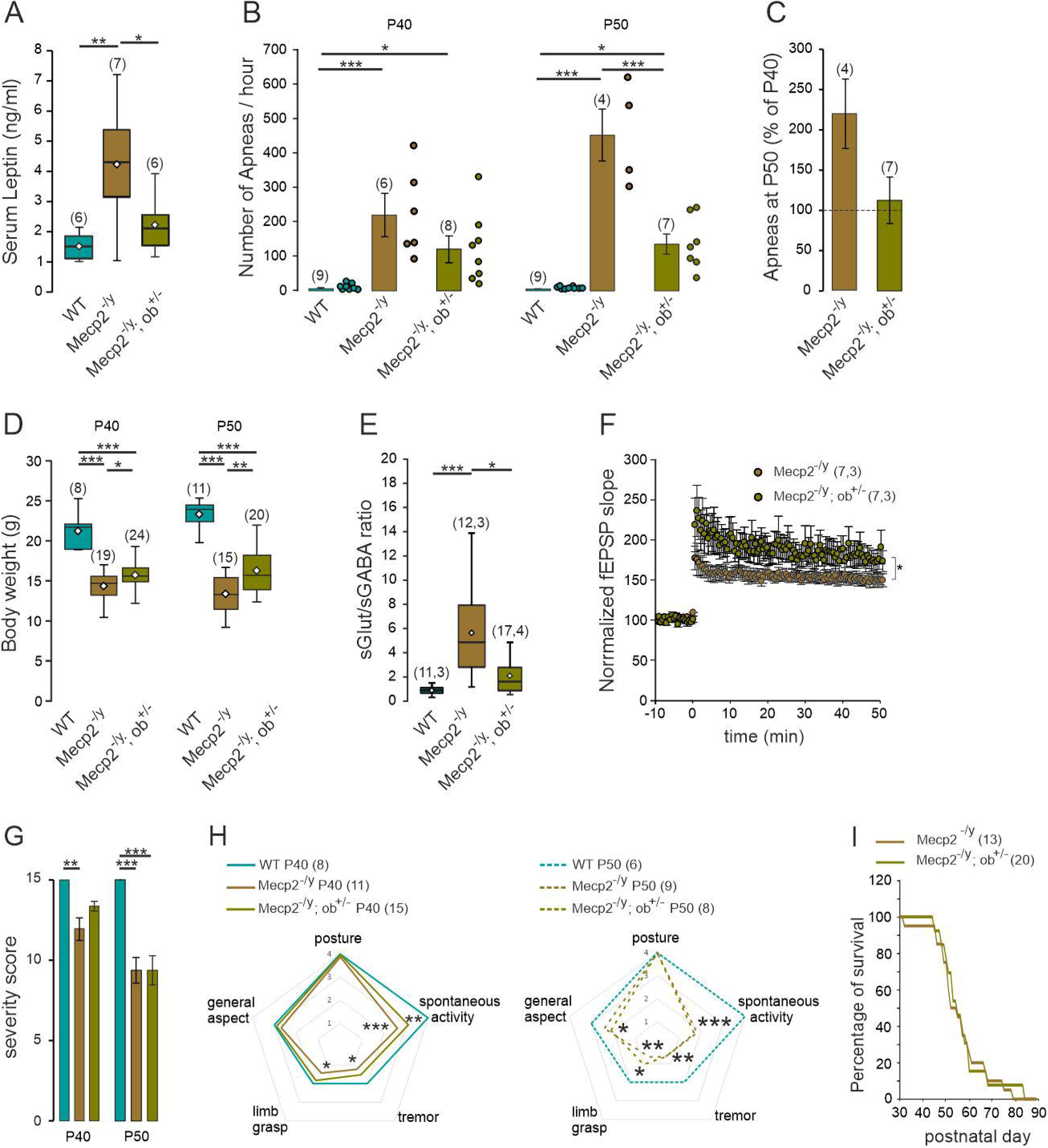
RTT-associated symptoms are improved in male leptin haplo insufficient mice. **A**) Box plot of serum leptin levels in P50 WT, *Mecp2*^-/y^ and *Mecp2*^-/y;ob/+^ littermates. **B**) Mean ±SEM plot of apnea frequency in P40 and P50 WT, *Mecp2*^-/y^ and *Mecp2*^-/y;ob/+^ littermates. Scatter plots show individual data points. **C**) Mean ±SEM change of apnea frequency at P50 (% of P40). **D**) Box plot of body weight in P40 and P50 WT, *Mecp2*^-/y^ and *Mecp2*^-/y;ob/+^ littermates. **E**) Box plots of the frequency ratio of spontaneous glutamatergic and GABAergic postsynaptic currents recorded from CA3 pyramidal neurons at P50. **F**) LTP at the Schaffer collateral-CA1 synapses of acute hippocampal slices from P50 *Mecp2*^-/y^ and *Mecp2*^-/y;ob/+^ mice. Numbers in parenthesis indicate the number of mice used and number of cells recorded. **G**) Average severity score of P40 and P50 WT, *Mecp2*^-/y^ and *Mecp2*^-/y;ob/+^ littermates. **H**) Radar plot of the symptoms scored in P40 and P50 WT, *Mecp2*^-/y^ and *Mecp2*^-/y;ob/+^ littermates. **I**) Plot of the percentage of surviving mice as a function of age. \**P* < 0.05; \*\**P* < 0.01; \*\*\**P* < 0.001. One-way ANOVA followed by a Tukey’s multiple comparison (A, B, D, E, G, H), Mann-Whitney test (C), two-tailed Kruskal-Wallis followed by a Dunn’s multiple comparison (H), repeated measure ANOVA (F).

Given that RTT primarily affects females, we wondered whether RTT symptoms were also alleviated in female *Mecp2^+/-^* mice that are haplo-insufficient for leptin (hereafter referred to as *Mecp2*^+/-^*;ob^+/-^* mice). A characteristic feature of the RTT mouse model on a C57BL/6J genetic background is that, unlike males, females gain weight as the disease progresses. The hyperleptinemia associated with this weight gain could therefore counterbalance haplo- insufficiency and prevent the normalization of leptin levels in *Mecp2*^+/-^*;ob^+/-^*mice. To address this point and identify a window for reliable assessment of leptin involvement in RTT symptoms, we performed a longitudinal measurement of circulating leptin levels in *Mecp2^+/-^, Mecp2*^+/-^*;ob^+/-^*and WT mice from postnatal day (P) 20 to 200. *Mecp2^+/-^* mice showed significantly elevated circulating leptin levels from P40 onwards compared to their WT littermates (**Figure 7A**). Moreover, at P40 and P50, but not at older stages, *Mecp2*^+/-;ob/+^ mice showed normalized leptin levels (**Figure 7A**). It is worth noting that from P100 *Mecp2^+/-^*and *Mecp2*^+/-^*;ob^+/-^* mice weighted more than WT (**Figure 7B**). This suggests that weight gain beyond P100, and the associated increase in leptin production, prevents normalization of leptin levels in *Mecp2*^+/-^*;ob^+/-^* mice.

**Figure 7:**
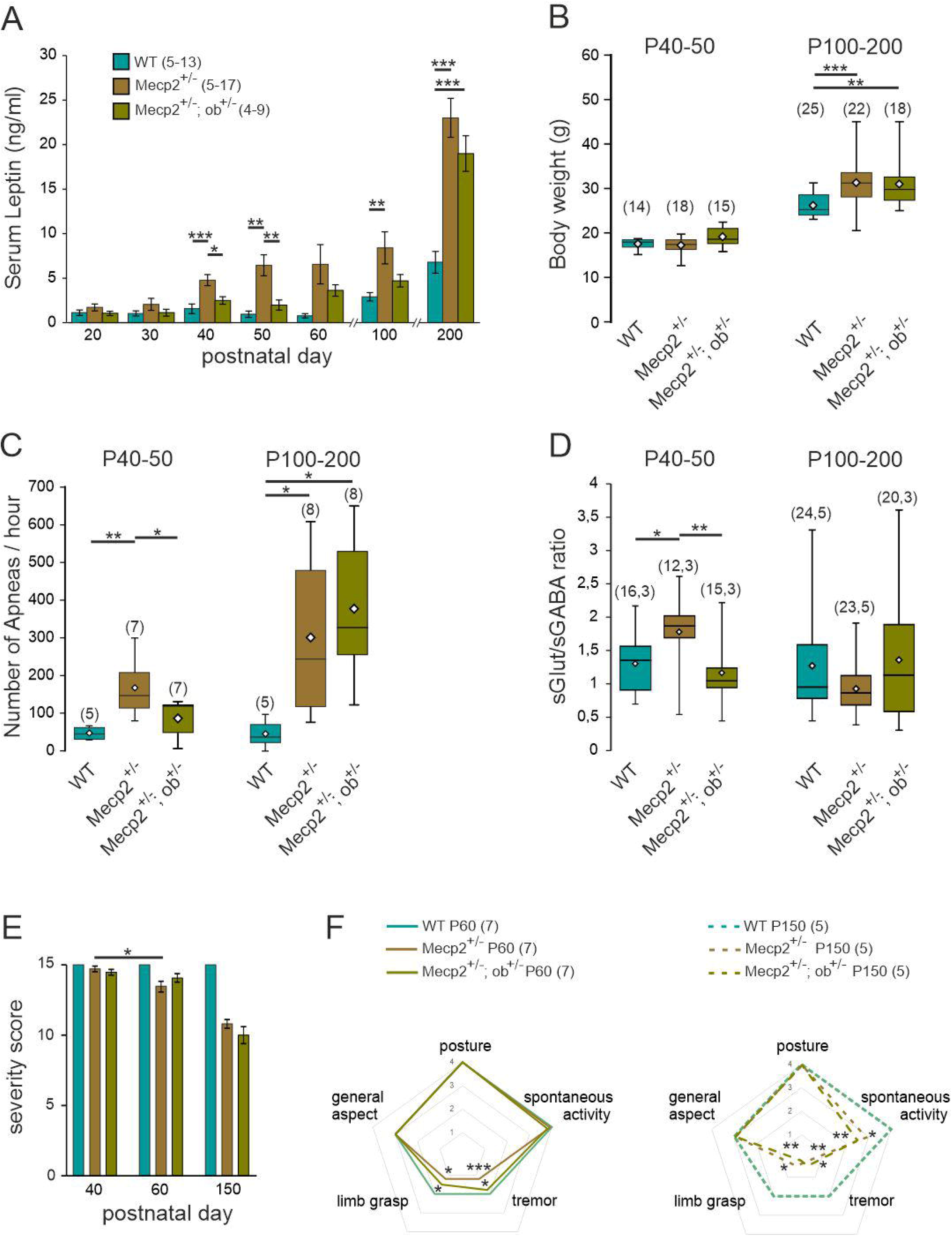
RTT-associated symptoms are improved in female leptin haplo insufficient mice. **A**) Mean ±SEM plot of serum leptin levels determined in blood samples taken at postnatal day (P) 20 every 10 days until P200, in female WT, *Mecp2*^+/-^ and *Mecp2*^+/-;ob/+^ littermates. **B-D**) Box plots of body weight (**B**), frequency of apneas (**C**) and frequency ratio of spontaneous glutamatergic and GABAergic postsynaptic currents (**D**) in P40-50 and P100-200 WT, *Mecp2*^+/-^ and *Mecp2*^+/-;ob/+^ littermates. **E**) Evolution of the severity score of WT, *Mecp2*^+/-^ and *Mecp2*^+/-;ob/+^ littermates. mean±SEM. **F**) Radar plot of the symptoms scored in P60 and P150 WT, *Mecp2*^+/-^ and *Mecp2*^+/-;ob/+^ littermates. \**P* < 0.05; \*\**P* < 0.01; \*\*\**P* < 0.001. Two-tailed Kruskal-Wallis followed by a Dunn’s multiple comparison (F), One-way ANOVA followed by a Tukey’s multiple comparison (A, B, C, D), Wilcoxon matched-pairs signed rank test (E).

Based on these observations, we conducted a behavioral and neuronal characterization of the female mice at two different stages: at an early stage (P40-50), when the leptin levels are normalized in *Mecp2*^+/-^*;ob^+/-^*mice and at a late stage (P100-200), when the leptin levels are comparable between *Mecp2^+/-^* and *Mecp2*^+/-^*;ob^+/-^* mice. We assessed several parameters including breathing, phenotypic score, E/I ratio, (**Figures 7C-F**). Notably, the *Mecp2*^+/-^*;ob^+/-^* mice showed an improvement in breathing activity at P40-50, but this improvement was not maintained at P100-200 (**Figure 7C**). Similarly, the E/I ratio recorded on CA3 hippocampal neurons was restored in P40-50 *Mecp2*^+/-^*;ob^+/-^* mice (**Figure 7D**). There was no significant difference in the E/I balance between WT, *Mecp2*^-/y^ and *Mecp2*^+/-^*;ob^+/-^* mice at P100-200 (**Figure 7D**). The longitudinal analysis of the health condition of the mice revealed a progressive decline in *Mecp^2-/y^* mice (**Figure 7E**), characterized by worsening of tremor and limb grasping from P50, followed by a decline in spontaneous activity from P100 (**Figure 7F**). In *Mecp2*^+/-^*;ob^+/-^* mice, the onset of tremors and limb grasping was delayed, though spontaneous activity showed no significant improvement (**Figure 7F**). Overall these data show that delaying the leptin rise delays the worsening of RTT-associated symptoms in *Mecp2*^+/-^*;ob^+/-^* mice.

## DISCUSSION

Here, we show that *Mecp2^-/y^* and *Mecp2^+/-^*mice have elevated circulating leptin levels compared to their age-matched WT littermates. Given the diverse central and peripheral roles of leptin, this finding provides a foundation for investigating its potential contribution to RTT pathogenesis. Remarkably, a 10-day treatment of male *Mecp2*-null mice from P40 with a leptin antagonist prevented the worsening of several symptoms including breathing difficulty, locomotor deficit, weight loss and degradation of general health condition. At the neuronal level, the anti-leptin treatment restored hippocampal synaptic plasticity as well as hippocampal and cortical excitatory/inhibitory balance. However, the leptin antagonism strategy showed limited effectiveness against anxiety, motor coordination deficits and early lethality. Consistent with the role of leptin to RTT pathogenesis, we also showed that leptin treatment of WT mice impaired the hippocampal excitatory/inhibitory balance, reduced hippocampal synaptic plasticity and caused breathing abnormality in male WT mice, thus mimicking some RTT symptoms. We further showed that genetic manipulation aimed at normalizing leptin level in male *Mecp2*-null mice prevents weight loss, the progression of breathing deficits, and rescues the excitatory/inhibitory balance and glutamatergic synaptic plasticity in the hippocampus. Finally, we showed that delaying the leptin rise in female *Mecp2*-deficient mice delays the worsening of their RTT-associated symptoms. These findings strengthen the link between leptin and RTT, providing a valuable insight into potential therapeutic strategy for this syndrome.

Elevated circulating leptin is consistently observed in patients with Rett syndrome ^7,8^ and in Mecp2-deficient mice ^9,10,28^. While chronic inflammation^37^, hormonal deregulations^38^, or medications ^39^ may contribute, increased adiposity alone is unlikely in patients, as hyperleptinemia occurs without changes in BMI or fat mass ^7,8,40,41^. In mice, however, elevated leptin is often associated with higher body weight^28^ ^42^ or fat mass ^9,10^. Notably, we found that symptomatic male *Mecp2*-null mice exhibit increased fat mass despite reduced body weight, together with elevated WAT leptin mRNA expression. The underlying mechanisms remain unclear, but they are likely to involve non-cell-autonomous processes. Supporting this hypothesis, previous studies showed that neuron-specific Mecp2 knockout mice (*Mecp2^tm1.1Jae^* KO mice) display both increased circulating leptin levels and higher WAT leptin mRNA expression ^9^, whereas adipocyte-specific Mecp2 knockout mice (*Mecp2^Adi^* KO mice) exhibit downregulated WAT leptin mRNA expression without changes in circulating leptin levels ^43^. Together, these findings suggest that both increased leptin synthesis per unit of adipose tissue and increased adiposity contribute to elevated circulating leptin. Finally, intramuscular adipose tissue, known to expand in neuromuscular and metabolic diseases ^44^, may also contribute, consistent with the muscle hypotrophy and fibrosis observed in symptomatic *Mecp2*-null mice ^45^.

In the present study, we used a pegylated super active mouse leptin antagonist (Peg-SMLA) that exhibits poor penetration in the central nervous system ^46^. The leptin antagonist inhibits leptin activity by two mechanisms: (i) by reducing the transport of circulating leptin across the blood brain barrier (BBB), thus reducing central leptin levels, and (ii) by blocking the binding of leptin to its peripheral receptors, thereby reducing peripheral leptin signaling. Similarly, haploinsufficiency for leptin mice reduces both central and peripheral actions of leptin. Several observations support the hypothesis that the beneficial effects of leptin antagonism rely on a reduction in the central action of leptin. Firstly, functional leptin receptors are expressed throughout the brain ^47^. *In vitro* studies indicate that leptin directly regulates the excitability and synaptic function of neurons within different brain regions, including the hippocampus ^15,23,24,26,48^ and nucleus tractus solitarius, which plays a critical role in the modulation of breathing, cardiovascular control and food intake ^49^. Furthermore, *in vivo* experiments demonstrate that local/systemic injections of leptin or chemogenetic/optogenetic activation of leptin receptor expressing neurons, modulate physiological and behavioral responses, and that targeted deletion of central leptin receptors prevent these effects ^50–56^. The respiratory network is complex and the effects of leptin on respiratory control are not yet fully understood. Nevertheless, alterations in the E/I balance have been reported in respiratory nuclei expressing leptin receptors, such as the nucleus tractus solitarius and the rostro ventrolateral medulla ^57,58^. The latter could contribute to apneustic and irregular breathing in *Mecp2*-deficient mice ^29^. More research is needed to determine whether and how leptin antagonism affects the E/I balance in these brain regions ^31^. Additionally, while our findings support the central effects of the treatments, we cannot completely rule out a peripheral effect, for example on carotid bodies. Further investigations are required to elucidate these peripheral effects in conjunction with the central mechanisms of leptin receptor antagonism.

Given the known facilitatory role of leptin on breathing and synaptic function, our present finding might initially appear counter-intuitive. However, it is worth noting that an excess of leptin can lead to a blunted response known as “leptin resistance”. Thus, rodents fed with a high-fat diet that causes elevated leptin levels and leptin resistance also exhibit altered hippocampal synaptic function, breathing, locomotor activity and anxiety level ^17,59^. Torres-Andrade and colleagues demonstrated that the absence of Mecp2 leads to hypothalamic leptin resistance in female mice ^28^. Consistent with this observation, we found that peripheral administration of leptin increases pSTAT3 immunostaining -a surrogate marker of leptin receptor activation-in hypothalamic nuclei of wild-type mice, but not in *Mecp2^-/y^* littermates (**supplementary Figure S4**). Furthermore, leptin injection in Mecp2^-/y^ mice has no effect on their weight, unlike in WT mice (**supplementary Figure S1D**). These findings support the notion that *Mecp2* deficiency disrupts leptin signaling, at least within hypothalamic circuits. It is conceivable that anti-leptin treatment or leptin haploinsufficiency, by lowering central leptin levels, alleviates leptin-resistance and thereby restore leptin sensitivity. This hypothesis is supported by studies in the context of obesity, where a reduction in circulating leptin levels has been shown to enhance leptin responsiveness, resulting in decreased food intake, increased energy expenditure, and improved glucose homeostasis ^60,61^. Conversely, daily leptin treatment of WT mice could lead to leptin resistance, which may be responsible for the symptoms observed in these mice. Interestingly, elevated leptin levels have also been observed in children with autism spectrum disorders ^62–64^. Yet, the clinical features seen in ASD only partially overlap with those of Rett syndrome, suggesting that the functional and physiological actions of leptin may depend, among other factors, on the specific mutations associated with the etiology of the disease.

We have shown that the anti-leptin treatment is well tolerated with no premature deaths or overt side effects, indicating it potential therapeutic application. Additionally, leptin antagonist therapy has been reported to be effective in preventing or treating diseases where elevated leptin levels are involved ^65,66^. It is noteworthy that leptin antagonist treatment has also shown improvements in bone mineralization ^67^ and muscle function ^68^, two peripheral symptoms observed in RTT. Furthermore, hyperleptinemia has been suggested to contribute to sympathetic overactivity and cardiac abnormalities observed in RTT ^8^. These observations collectively suggest that leptin may play a role in various central and peripheral RTT clinical manifestations underscoring the broad therapeutic potential of this approach.

Given the progressive nature of Rett symptoms and the high inter-individual variability, we assessed their evolution (i.e., the difference between the values at the beginning and at the end of treatment) to estimate the potential efficacy of the leptin antagonist treatment. Our data indicate that treatment with the leptin antagonist during the symptomatic period delays the progression of certain symptoms (i.e. breathing, locomotor activity, weight loss, general health status) but does not lead to a complete recovery. The notion of a pre-symptomatic period in RTT has recently been questioned, as studies have shown subtle symptoms in RTT girls and early alterations in animal models before the appearance of overt symptoms ^69^. Therefore, while translational applications may pose challenges, it will be interesting in the future to determine whether “pre-symptomatic” treatments have more significant positive effects. Such interventions could shed light on potential therapeutic strategies for intervening at an earlier stage to possibly improve treatment outcomes in RTT.

## MATERIALS and METHODS

### Animals

All animal procedures were carried out in accordance with the European Union Directive of 22 September (2010/63/EU) and have been approved by the Ethical Committee for Animal Experimentation (APAFIS-Numbers 17605-2018-1119-1115-7094 and 41156-2023-0220-1011-9732) delivered by the French Ministry of Education and Research.

Experiments were performed on male wild type and *Mecp^2tm1-1Bird^*mice ^70^ C57BL/6J genetic background (Jackson Laboratory, stock number 003890). Hemizygous mutant males (hereafter referred as to *Mepc2^-/y^* mice) were generated by crossing heterozygous females (*Mecp2*^+/-^) with C57BL/6 wild type males. *Mecp2*^-/y^ and *Mecp2*^+/-^ mice haplo insufficient for leptin (hereafter referred as to *Mecp2*^-/y^*;ob^+/-^*and *Mecp2*^+/-^*;ob^+/-^*) were generated by crossing heterozygous females *Mecp2*^+/-^ with C57BL/6 heterozygous leptin deficient males mice (Jackson Laboratory genotyping protocol, strains B6.Cg-Lepob/J, ID 000632). Animals were housed in a temperature-controlled environment with a 12h light/dark cycle and free access to food and water. Genotyping was performed by PCR techniques according to Jackson Laboratory protocols.

### Leptin and anti-leptin injection

Recombinant murine leptin (Peprotech, New Jersey, USA) was reconstituted in sterile water and daily injected (5 µg/g) sub-cutaneous at 12-14 hr a.m during 10 days starting at postnatal day (P) 40. The pegylated super active mouse leptin antagonist (Peg-SMLA, Protein Laboratories, Rehovot, Israel) was reconstituted in sterile water ^46^. As the half-life of the Peg-SMLA was about 24 hours, mice were injected (5 µg/g) sub-cutaneous at 12-14 hr a.m every other day during 10 days starting at P40. Control mice received same volume injections of vehicle.

### Leptin and BDNF immunoassay

Blood samples were collected by two routes: submandibular bleeding, performed every 10 days from postnatal day 20 to 90 in female mice or decapitation (trunk blood samples) at P15, 30 and 50 in male mice and at P100-200 in female mice. Blood samples were centrifuged (10.000 rpm, 10 min, 4°C) immediately after collection at 10–11 hr a.m. Quantification of endogenous leptin was performed with Mouse Leptin ELISA Kit (BioVendor R&D^R^, Brno, Czech Republic) following the manufacturer’s protocol. The measured concentration of samples was calculated from the standard curve and expressed as ng/ml.

Hippocampal tissues were homogenized in RIPA buffer (150 mM NaCl, 1% Triton X100, 0.1% SDS, 50 mM Tris HCl), pH 8, containing protease inhibitors (Complete Mini; Roche). Lysates were centrifuged (10.000 g for 30 min at 4 °C). Loading was 200 µg of proteins as determined using a modified Bradford reaction (BioRad Laboratories). Quantification of mature- and proBDNF was performed with both proBDNF and mature BDNF Rapid ELISA Kit (Biosensis Pty Ltd., Thebarton, SA, Australia) in the concentrated solutions following the manufacturer’s protocol.

### Real-time reverse transcription quantitative polymerase chain reaction

Visceral and inguinal fat pads and gastrocnemius muscles were dissected, rapidly frozen in liquid nitrogen and stored at -80°C. Total RNAs from fat pads and gastrocnemius muscle were isolated using a RNeasy Mini kit (74106, Qiagen, Germany) and RNA Fibrous Tissue Mini kit (74704, Qiagen) respectively, following the manufacturer’s instructions and quantified by reading the absorbance at 260 nm (NanoPhotometer, IMPLEN). RNAs were converted to cDNA using 1 µg RNA and a QuantiTect Reverse Transcription kit (Qiagen) according to manufacturer’s instructions. Quantitative RT-PCR was performed with a Light Cycler 480 SYBR Green IMaster (Roche Applied Science) following the manufacturer’s instructions, using the following oligonucleotides (QuantiTect Primer Assays, Qiagen): Leptin (Mm_Lep_1_SG QT00164360 (NM_008493)), and glyceraldehyde-3-phosphate dehydrogenase (GAPDH, QT001199633). Relative mRNA values were calculated using the LC480 software and GAPDH as the housekeeping gene. PCR was performed in triplicate.

### Slice preparation and electrophysiological recordings

Brains were removed and immersed into ice-cold (2° to 4°C) artificial cerebrospinal fluid (ACSF) with the following composition: 126 mM NaCl, 3.5 mM KCl, 2 mM CaCl2, 1.3 mM MgCl2, 1.2 mM NaH2PO4, 25 mM NaHCO3, and 11 mM glucose (pH 7.4) equilibrated with 95% O2 and 5% CO2. Hippocampal slices (350µm thick) were cut using a vibrating microtome (Leica VT1000S, Germany) in ice-cold oxygenated choline-replaced ACSF and were allowed to recover at least 90 min in ACSF at room temperature (25°C). Slices were then transferred to a submerged recording chamber perfused with oxygenated (95% O2 and 5% CO2) ACSF (3 ml/min) at 34°C.

*Whole cell recordings* were performed from hippocampal CA3 and somatosensory layer V pyramidal neurons in the voltage-clamp mode using an axopatch 200B amplifier (Molecular Devices LLC, San Jose, USA). To assess the excitatory/inhibitory balance, the glass recording electrodes (4-7 MΩ) were filled with a solution containing 100 mM K-gluconate, 13 mM KCl, 10 mM Hepes, 1.1 mM EGTA, 0.1 mM CaCl2, 4 mM Mg–adenosine 5′-triphosphate, and 0.3 mM Na–guanosine 5′-triphosphate. The pH of the intracellular solution was adjusted to 7.2, and the osmolality was adjusted to 280 mOsmol liter−1. With this solution, GABA_A_-receptor mediated postsynaptic currents (GABA_A_-PSCs) reversed at −70 mV. GABA_A_-PSCs and Glut-PSCs were recorded at a holding potential of −45 and -70 mV respectively. Spontaneous synaptic currents, were recorded with Axoscope software version 8.1 (Molecular Devices LLC, San Jose, USA) and analyzed offline with Mini Analysis Program version 6.0 (Synaptosoft).

*Field recordings* were performed in area CA1. A cut was made between CA1 and CA3. The Schaffer collaterals/commissural fibers were stimulated using a bipolar tungsten electrode (66 μm; A-M Systems, WA, USA) placed on the surface of the stratum radiatum of CA1 (10–50 μs, 5–15 V, 0.03 Hz). Extracellular tungsten electrodes (California Fine Wire,CA, USA) were used to record dendritic field excitatory postsynaptic potentials (fEPSP) from the stratum radiatum of the CA1 region. The signals were amplified using a DAM80 amplifier (WPI, UK), digitized with an Axon Digidata 1550B (Molecular Devices, CA, USA), recorded with Axoscope software version 8.1 (Molecular Devices, CA, USA) and analyzed offline with Mini Analysis Program version 6.0 (Synaptosoft, GA, USA) by measuring the onset (a 30–70% rising phase) slope of the fEPSP. LTP was induced by three 100 Hz trains of 100 stimuli 30 sec apart at 50% of the maximal intensity. The slope of the fEPSP was measured and expressed relative to the preconditioning baseline.

### pSTAT3 Immunohistochemistry

Mice were deeply anesthetized with tiletamine/zolazepam (Zoletil, 40 mg/kg) and medetomidine (Domitor, 0.6 mg/kg) and trans-cardially perfused with 4% paraformaldehyde (PFA) in 0.1 M phosphate buffer (pH 7.4). After perfusion, brains were removed and post-fixed through immersion in cold solution of 4% PFA for 24 h and then stored at 4°C in Phosphate-Buffered Saline (PBS) with **0.01%** sodium azide until processing. Serial coronal brain sections (40 µm thickness) at the hypothalamic level (from the bregma coordinates -0.58 to -2.18) were done using a vibratome (Leica VT1000S, Germany) and collected in PBS.

Three randomly chosen sections per animal of the same coordinates range for each hypothalamic nucleus examined were used for the immuhistochemistry processing (bregma -0.80 to -1.00 for the paraventricular nucleus of the hypothalamus (PVN); -1.06 to -1.34 for the lateral hypothalamic area (LH); -1.46 to -1.70 for the dorsomedial hypothalamic nu (DM); -1.46 to -1.82 for the ventromedial hypothalamic nu (VMH); -1.58 to -1.94 for the arcuate hypothalamic nu (Arc)). The free-floating sections were first rinsed in Tris buffer salin (TBS) and pre-treated in 0.3% Glycine for 30 min at room temperature (RT). They were then permeabilized for 30 min in 0.3% triton x100 and were blocked in 5% normal donkey serum and 5% bovine serum albumin (BSA) for 1 h at RT. Section were incubated overnight at 4°C in pSTAT3 antibody (1/200 Cat 9131, cell Signaling) diluted in TBS with 1% BSA. On the next day, sections were washed four times in TBS and incubated with a donkey anti rabbit Alexa Fluor 568 labeled secondary antibody (1/500) for 2 h. The stained sections were mounted on coverslips with mounting reagent (FluorSave reagent, Calbiochem) and kept away from light until analysis.

The immunostaining in the nuclei of interest was acquired at 20x magnification using a Zeiss AxioImager M2 Apo1.2 microscope from IBDM imaging facility. Each hypothalamic structure examined was delineated and the nuclear profiles of all pSTAT3 positive labeled cells were manually counted from 3 sections using Fiji Image J analysis software from the NIH, to exclude any bias that could result from changes in cell soma size or shape among experimental conditions. The average number of cells counted in the 3 sections per area in each mouse was taken for statistical comparisons.

### Plethysmography

The breathing activity of non-anesthetized freely moving mice was recorded using a constant flow whole body plethysmograph (EMKA technologies, Paris, France) with 200ml animal chambers maintained at 25±0.5 ^◦^C and ventilated with air (600ml min^-1^). The breathing activity of pairs of WT and *Mecp2*^-/y^ mice littermates was simultaneously recorded. Mice were allowed to acclimate to the experimental room for 1 hour and to the plethysmography chamber and airflow for approximatively 30 mins before breathing measurement. Breathing activity was recorded during 1 hour. A differential pressure transducer measured the changes in pressure in the body chamber resulting from the animal’s respiration. Signals from the pressure transducer were stored on a personal computer and analyzed offline via the Spike 2 interface and software (Cambridge Electronic Design, Cambridge, UK). Only periods of quiet breathing without body movements were analyzed, during which the number of apneas (> three respiratory cycles) per hour was quantified.

### General health scoring and lifespan

Weight was measured every day from the beginning of the treatment. The phenotypic score was evaluated the day before treatment and the last day of treatment. Severity score, typically used in RTT phenotypic assessments ^70^, was used to group animals into 4 severity classes: absence of phenotype (4) to severe phenotype (0). The parameters scored are: tremor, posture, limb grasp, spontaneous activity in the home cage and general aspect. To be noticed, mice that rapidly lost weight were euthanized for ethical reasons. The day of the sacrifice was considered as the endpoint of lifespan assessment.

### Accelerating rotarod

A rotarod apparatus (Biological Research Apparatus, Gemonio, Italy) was used to measure the motor coordination. After a 5 mins habituation session (4 r.p.m), each mouse was given three trials with the rate of rotation increasing from 4 to 40 r.p.m. over 5 mins. The trial ended when the mouse fell from the rod or remained on the rotarod for at least 5 mins. The time spent on the rotarod was recorded by an automated unit, which stopped as the mouse fell. The mouse was placed back in its home cage for 10 mins between each trial. The latency to fall was determined as the best of the 3 trials.

### Behavior

All the behavioral tests were performed by Phenotype Expertise, Inc. (France). For all tests, animals were first acclimated to the behavioral room for 30 minutes. WT and *Mecp2*^-/y^ mice underwent elevated plus maze, open field, three chamber test and spontaneous social interaction test at P50. WT and *Mecp2*^-/y^ treated mice underwent elevated plus maze and open field at P40, before the beginning of the treatment, and at P50, at the end of the treatment.

#### Elevated-Plus Maze

The EPM was used to assess anxiety state of animals. The device consists of a labyrinth of 4 arms 5 cm wide located 80 cm above the ground. Two opposite arms are open (without wall) while the other two arms are closed by side walls. The light intensity was adjusted to 20 Lux on the open arms. Mice were initially placed on the central platform and left free to explore the cross-shaped labyrinth for 5 minutes. Maze was cleaned and wiped with H2O and with 70% ethanol between each mouse. Animal movement was video-tracked using Ethovision software 11.5 (Noldus). Time spent in open and closed arms, the number of entries in open arms, as well as the distance covered, are directly measured by the software.

#### Open-field

Open field was used to evaluate both the locomotor activity of the animals. Mice were individually placed in a 40 x 40 cm square arena with an indirect illumination of 60 lux. Mouse movement was video-tracked using Ethovision software 11.5 (Noldus) for 10 minutes. Total distance traveled and time in center (exclusion of a 5 cm border arena) are directly measured by the software. Grooming (time and events) and rearing were manually counted in live using manual functions of the software, by an experimented behaviorist. The open-field arena was cleaned and wiped with H2O and with 70% ethanol between each mouse.

#### Three-chamber social preference test

A square arena 40 x 40 cm was used, with the wired cages placed in opposite diagonal corners. The task was carried out in four trials. The three-chambers apparatus was cleaned and wiped with water and with 70% ethanol between each trial and each mouse. In the first trial (habituation), a test mouse was placed in the center of the arena and was allowed to freely explore each chamber. The mouse was video-tracked for 5 min using Ethovision software. At the end of the trial, the animal was briefly removed from the arena to allow for the preparation of the following trial. In the second trial (social exploration), a 8-weeks old C57BL/6J congener mouse (S1) was placed randomly in one of the two wire cages to avoid a place preference. The second wire cage remained empty (E). Then the test mouse was placed in the center of the arena and allowed to freely explore for 10 min. A second 8-weeks old C57BL/6J congener mouse (S2) was placed in the second wire cage for the third trial (social discrimination). Thus, the tested mouse had the choice between a familiar mouse (S1) and a new stranger mouse (S2) for 10 min. At the end of the trial, the mouse was returned to home-cage for 30 min. For the fourth trial (short-term social memory), S2 was replaced by a new stranger mouse (S3), the familiar mouse (S1) staying the same. Then tested mouse was allowed to freely explore the arena for 10 min. Time spent in each chamber and time of contact with each wire cage (with a mouse or empty) were calculated using Ethovision software. The measure of the real social contact is represented by the time spent in nose-to-nose interactions with the unfamiliar or familiar mouse. This test was performed using grouped house mice of 4 months old.

#### Spontaneous social interaction test

The tested mouse was placed into the same OF arenas that were used for the OF test and allowed to explore this empty arena for 2.5 min. Immediately following this initial stage, the mouse was again allowed to explore the OF for an additional 10 min with the target (Swiss mouse) present. Mouse movement was video-tracked using Ethovision software 11.5 (Noldus) and the time spent in nose-to-nose, nose-to-body, and nose-to-genitals interactions was measured.

### Statistics

Statistical analyses were conducted with GraphPad Prism (GraphPad software 5.01). Shapiro-Wilk normality test was used to determine the normality of distributions. We used a Two-tailed Mann-Whitney *U* test or Two-tailed unpaired *t*-test for comparison between two independent groups, a Wilcoxon matched pairs signed rank test or Two-tailed paired *t*-test for comparison between two matched data, a Two-tailed Kruskal-Wallis test followed by a Dunn’s multiple comparison or a two-way ANOVA followed by a Tukey’s multiple comparison for comparison between three or more independent groups, and a Fisher exact test for nominal comparison between two independent groups. The effect of tetanic stimulation of the slope of field EPSPs was analyzed using repeated measure ANOVA. The survival analysis was performed using a Kaplan-Meier log-rank test. Possible outliers within an experimental group were identified with Grubb’s test and discarded from the final analysis. To ensure the consistency and reproducibility of our results, we conducted repeated trials in different acute hippocampal slices prepared from at least three different animals from three different littermates. All data are expressed as mean ± standard error to the mean (S.E.M.). For results expressed as percent of WT (i.e. serum and mRNA leptin, Figure 1, all values (WT and *Mecp2^-/y^* mice) were normalized to the mean WT value. In the figures, box plots represent the 1^rst^ and 3^rd^ quartiles; whiskers show data range; horizontal lines show the median.

## ACKNOWLEDGMENTS

We thank the members of the Molecular and Cellular Biology facility, Histological facility (Marie Kurz) and Animal facility (Quenol Cesar) at INMED. We thank Dr. L. Kerkerian-Le Goff for critical reading of the manuscript.

## DATA AVAILABILITY

All datasets generated for this study are available on request to the corresponding author.

## AUTHOR CONTRIBUTIONS

J-LG, YB, CM and GAW conceived the experiments and wrote the manuscript. YB, J-LG, DD, MB-H, PS and VV performed the experiments. CS performed the behavioral analysis. J-CG bred the colony. All authors approved the final version of the manuscript.

---

**Supplementary Fig S1:** Leptin treatment has no effect on *Mecp2^-/y^* mice. P40 *Mecp2^-/y^* received daily sub-cutaneous injection of leptin recombinant (5µg/g) during 10 days. Sham mice received the same volume of vehicle. **A-C**) Box plots of the percentage of change (% of P40) of apnea frequency (**A**), breathing irregularity score (**B**) and minute ventilation (**C**) in sham- and treated- *Mecp2*^-/y^ mice. **D**) Body weight change (% of P40) as a function of age in sham-anti-leptin treated *Mecp2^-/y^*mice. **E)** Box plots of the frequency ratio of spontaneous glutamatergic and GABAergic postsynaptic currents recorded on CA3 pyramidal neurons of sham- and leptin treated- *Mecp2*^-/y^ mice at P50. Numbers in parenthesis indicate the number of mice used. Two-tailed unpaired t-test.

**Supplementary Fig S2:** Lack of differences in social and cognitive behavior between P50 WT and *Mecp2^-/y^*mice. **A, B**) Mean ± SEM plots of the time interacting of P50 WT (A) and *Mecp2^-/y^* (B) mice during the different phase of the 3 chambers test. **C**) Box plots of the time interacting of P50 WT and *Mecp2^-/y^* mice with a stranger mouse in the spontaneous social interaction test. **D**) Mean ± SEM plots of the time interacting and index preference of P50 WT and *Mecp2^-/y^*mice in the novel object recognition test. Numbers in parenthesis indicate the number of mice used. \*\**P* < 0.01; \*\*\**P* < 0.001, two-tailed paired *t*-test (A, B, D), two-tailed unpaired *t*-test (B, D).

**Supplementary Fig S3:** mature and pro-brain derived neurotrophic factor (BDNF) proteins expression in WT and *Mecp2^-/y^* mice. **A-C**) Box plots of mature (A) and pro (B) BDNF protein expressions in hippocampal samples taken from P50 WT and *Mecp2^-/y^* mice, using ELISA kit. P40 WT and *Mecp2^-/y^* received sub-cutaneous injection of vehicle (sham) or anti-leptin (5µg/g), every other day, during 10 days. \**P* < 0.05, One-way ANOVA followed by a Tukey’s multiple comparison.

**Supplementary Fig S4:** Lack of phenotypic differences between WT and *Mecp2^+/-^* mice. **B**) Representative images of pSTAT3 labeling within the arcuate nucleus (Arc) the lateral hypothalamic area (LHA) the ventromedial hypothalamus (VMH) and the dorsomedial hypothalamus (DM) of leptin-treated and sham *Mecp2^-/y^*mice (B) (bregma level -1.94). Note that a high number of pSTAT3 immuno-positive cells, intensely labeled, is detected in the DM and the Arc in both conditions. However, the significant but less intense pSTAT3 labeled cells observed in the VMH of sham *Mecp2^-/y^*mice is no longer observed in leptin-treated *Mecp2^-/y^* mice. 3V: third ventricle; Opt: optic tract; fr: fornix; delineated STN: Subthalamic nucleus; delineated cp: cerebral peduncle, Scale bars, 250 µm. **B**) Percentage of increase of pSTAT3 positive cells induced by sub-cutaneous injection of leptin in WT and *Mecp2*^-/y^ mice. LH: Lateral hypothalamic area; DM: dorsomedial nucleus; VMH: ventromedial nucleus; Arc: arcuate nucleus; PVN: periventricular nucleus.

## REFERENCES

1. Amir, R. E. et al. Rett syndrome is caused by mutations in X-linked MECP2, encoding methyl-CpG-binding protein 2. Nat Genet 23, 185–188 (1999).

2. Cheval, H. et al. Postnatal inactivation reveals enhanced requirement for MeCP2 at distinct age windows. Hum Mol Genet 21, 3806–3814 (2012).

3. Sandweiss, A. J., Brandt, V. L. & Zoghbi, H. Y. Advances in understanding of Rett syndrome and MECP2 duplication syndrome: prospects for future therapies. Lancet Neurol 19, 689–698 (2020).

4. Ricceri, L., De Filippis, B. & Laviola, G. Mouse models of Rett syndrome: from behavioural phenotyping to preclinical evaluation of new therapeutic approaches. Behav Pharmacol 19, 501–517 (2008).

5. Guy, J., Gan, J., Selfridge, J., Cobb, S. & Bird, A. Reversal of neurological defects in a mouse model of Rett syndrome. Science 315, 1143–1147 (2007).

6. Ehinger, Y., Matagne, V., Villard, L. & Roux, J.-C. Rett syndrome from bench to bedside: recent advances. F1000Res 7, 398 (2018).

7. Blardi, P. et al. Rett syndrome and plasma leptin levels. J Pediatr 150, 37–39 (2007).

8. Acampa, M. et al. Sympathetic overactivity and plasma leptin levels in Rett syndrome. Neurosci. Lett. 432, 69–72 (2008).

9. Park, M. J. et al. Anaplerotic triheptanoin diet enhances mitochondrial substrate use to remodel the metabolome and improve lifespan, motor function, and sociability in MeCP2-null mice. PLoS One 9, e109527 (2014).

10. Fukuhara, S. et al. High-fat diet accelerates extreme obesity with hyperphagia in female heterozygous Mecp2-null mice. PLoS One 14, e0210184 (2019).

11. Ahima, R. S. & Flier, J. S. Leptin. Annual Review of Physiology 62, 413–437 (2000).

12. Davis, J. F., Choi, D. L. & Benoit, S. C. Insulin, leptin and reward. Trends Endocrinol Metab 21, 68–74 (2010).

13. Harvey, J. Food for Thought: Leptin and Hippocampal Synaptic Function. Front Pharmacol 13, 882158 (2022).

14. Guo, M. et al. Forebrain glutamatergic neurons mediate leptin action on depression-like behaviors and synaptic depression. Transl Psychiatry 2, e83 (2012).

15. Shanley, L. J., Irving, A. J., Rae, M. G., Ashford, M. L. J. & Harvey, J. Leptin inhibits rat hippocampal neurons via activation of large conductance calcium-activated K+ channels. Nat Neurosci 5, 299–300 (2002).

16. Mora-Muñoz, L., Guerrero-Naranjo, A., Rodríguez-Jimenez, E. A., Mastronardi, C. A. & Velez-van-Meerbeke, A. Leptin: role over central nervous system in epilepsy. BMC Neurosci 19, 51 (2018).

17. Gauda, E. B. et al. Leptin: Master Regulator of Biological Functions that Affects Breathing. Compr Physiol 10, 1047–1083 (2020).

18. Huo, L. et al. Leptin-dependent control of glucose balance and locomotor activity by POMC neurons. Cell Metab 9, 537–547 (2009).

19. Signore, A. P., Zhang, F., Weng, Z., Gao, Y. & Chen, J. Leptin neuroprotection in the CNS: mechanisms and therapeutic potentials. J. Neurochem. 106, 1977–1990 (2008).

20. Lim, G., Wang, S., Zhang, Y., Tian, Y. & Mao, J. Spinal leptin contributes to the pathogenesis of neuropathic pain in rodents. J Clin Invest 119, 295–304 (2009).

21. Bouret, S. G., Draper, S. J. & Simerly, R. B. Trophic action of leptin on hypothalamic neurons that regulate feeding. Science 304, 108–110 (2004).

22. Moult, P. R. & Harvey, J. Hormonal regulation of hippocampal dendritic morphology and synaptic plasticity. Cell Adh Migr 2, 269–275 (2008).

23. Dhar, M. et al. Leptin-induced spine formation requires TrpC channels and the CaM kinase cascade in the hippocampus. J. Neurosci. 34, 10022–10033 (2014).

24. Dumon, C. et al. The adipocyte hormone leptin sets the emergence of hippocampal inhibition in mice. Elife 7, e36726 (2018).

25. Sahin, G. S. et al. Leptin stimulates synaptogenesis in hippocampal neurons via KLF4 and SOCS3 inhibition of STAT3 signaling. Mol Cell Neurosci 106, 103500 (2020).

26. Sahin, G. S. et al. Leptin increases GABAergic synaptogenesis through the Rho guanine exchange factor β-PIX in developing hippocampal neurons. Sci Signal 14, eabe4111 (2021).

27. Harvey, J. Leptin: a multifaceted hormone in the central nervous system. Mol Neurobiol 28, 245–258 (2003).

28. Torres-Andrade, R. et al. The increase in body weight induced by lack of methyl CpG binding protein-2 is associated with altered leptin signalling in the hypothalamus. Exp. Physiol. 99, 1229–1240 (2014).

29. Ramirez, J.-M., Ward, C. S. & Neul, J. L. Breathing challenges in Rett syndrome: lessons learned from humans and animal models. Respir Physiol Neurobiol 189, 280–287 (2013).

30. Katz, D. M., Dutschmann, M., Ramirez, J.-M. & Hilaire, G. Breathing disorders in Rett syndrome: progressive neurochemical dysfunction in the respiratory network after birth. Respir Physiol Neurobiol 168, 101–108 (2009).

31. Li, W. Excitation and Inhibition Imbalance in Rett Syndrome. Front Neurosci 16, 825063 (2022).

32. Calfa, G., Li, W., Rutherford, J. M. & Pozzo-Miller, L. Excitation/inhibition imbalance and impaired synaptic inhibition in hippocampal area CA3 of Mecp2 knockout mice. Hippocampus 25, 159–168 (2015).

33. Asaka, Y., Jugloff, D. G. M., Zhang, L., Eubanks, J. H. & Fitzsimonds, R. M. Hippocampal synaptic plasticity is impaired in the Mecp2-null mouse model of Rett syndrome. Neurobiol Dis 21, 217–227 (2006).

34. Ehinger, Y. et al. Huntingtin phosphorylation governs BDNF homeostasis and improves the phenotype of Mecp2 knockout mice. EMBO Mol Med 12, e10889 (2020).

35. Liao, G.-Y. et al. Dendritically targeted Bdnf mRNA is essential for energy balance and response to leptin. Nat Med 18, 564–571 (2012).

36. Yamada, N. et al. Impaired CNS leptin action is implicated in depression associated with obesity. Endocrinology 152, 2634–2643 (2011).

37. Cortelazzo, A. et al. Subclinical inflammatory status in Rett syndrome. Mediators Inflamm 2014, 480980 (2014).

38. Stagi, S. et al. Thyroid function in Rett syndrome. Horm Res Paediatr 83, 118–125 (2015).

39. Belcastro, V., D’Egidio, C., Striano, P. & Verrotti, A. Metabolic and endocrine effects of valproic acid chronic treatment. Epilepsy Res 107, 1–8 (2013).

40. Motil, K. J., Schultz, R., Brown, B., Glaze, D. G. & Percy, A. K. Altered energy balance may account for growth failure in Rett syndrome. J Child Neurol 9, 315–319 (1994).

41. Motil, K. J., Schultz, R. J., Wong, W. W. & Glaze, D. G. Increased energy expenditure associated with repetitive involuntary movement does not contribute to growth failure in girls with Rett syndrome. J Pediatr 132, 228–233 (1998).

42. Wang, X., Lacza, Z., Sun, Y. E. & Han, W. Leptin resistance and obesity in mice with deletion of methyl-CpG-binding protein 2 (MeCP2) in hypothalamic pro-opiomelanocortin (POMC) neurons. Diabetologia 57, 236–245 (2014).

43. Liu, C. et al. Fat-Specific Knockout of Mecp2 Upregulates Slpi to Reduce Obesity by Enhancing Browning. Diabetes 69, 35–47 (2019).

44. Sciorati, C., Clementi, E., Manfredi, A. A. & Rovere-Querini, P. Fat deposition and accumulation in the damaged and inflamed skeletal muscle: cellular and molecular players. Cell Mol Life Sci 72, 2135–2156 (2015).

45. Conti, V. et al. MeCP2 Affects Skeletal Muscle Growth and Morphology through Non Cell-Autonomous Mechanisms. PLoS One 10, e0130183 (2015).

46. Elinav, E. et al. Pegylated leptin antagonist is a potent orexigenic agent: preparation and mechanism of activity. Endocrinology 150, 3083–3091 (2009).

47. Caron, E., Sachot, C., Prevot, V. & Bouret, S. G. Distribution of leptin-sensitive cells in the postnatal and adult mouse brain. J. Comp. Neurol. 518, 459–476 (2010).

48. Guimond, D. et al. Leptin potentiates GABAergic synaptic transmission in the developing rodent hippocampus. Frontiers in Cellular Neuroscience 8, 235 (2014).

49. Neyens, D. et al. Leptin Sensitizes NTS Neurons to Vagal Input by Increasing Postsynaptic NMDA Receptor Currents. J Neurosci 40, 7054–7064 (2020).

50. Oomura, Y. et al. Leptin facilitates learning and memory performance and enhances hippocampal CA1 long-term potentiation and CaMK II phosphorylation in rats. Peptides 27, 2738–2749 (2006).

51. Inyushkin, A. N., Inyushkina, E. M. & Merkulova, N. A. Respiratory responses to microinjections of leptin into the solitary tract nucleus. Neurosci Behav Physiol 39, 231–240 (2009).

52. Vong, L. et al. Leptin action on GABAergic neurons prevents obesity and reduces inhibitory tone to POMC neurons. Neuron 71, 142–154 (2011).

53. Guo, M., Huang, T.-Y., Garza, J. C., Chua, S. C. & Lu, X.-Y. Selective deletion of leptin receptors in adult hippocampus induces depression-related behaviours. Int. J. Neuropsychopharmacol. 16, 857–867 (2013).

54. Zuure, W. A., Roberts, A. L., Quennell, J. H. & Anderson, G. M. Leptin signaling in GABA neurons, but not glutamate neurons, is required for reproductive function. J Neurosci 33, 17874–17883 (2013).

55. Bassi, M. et al. Leptin into the ventrolateral medulla facilitates chemorespiratory response in leptin-deficient (ob/ob) mice. Acta Physiol (Oxf) 211, 240–248 (2014).

56. Bassi, M. et al. Activation of the brain melanocortin system is required for leptin-induced modulation of chemorespiratory function. Acta Physiol (Oxf) 213, 893–901 (2015).

57. Medrihan, L. et al. Early defects of GABAergic synapses in the brain stem of a MeCP2 mouse model of Rett syndrome. J Neurophysiol 99, 112–121 (2008).

58. Chen, C.-Y. et al. Defective GABAergic neurotransmission in the nucleus tractus solitarius in Mecp2-null mice, a model of Rett syndrome. Neurobiol Dis 109, 25–32 (2018).

59. Farr, O. M., Tsoukas, M. A. & Mantzoros, C. S. Leptin and the brain: influences on brain development, cognitive functioning and psychiatric disorders. Metabolism 64, 114–130 (2015).

60. Zhao, S. et al. Partial Leptin Reduction as an Insulin Sensitization and Weight Loss Strategy. Cell Metab 30, 706–719.e6 (2019).

61. Zhao, S. et al. Partial leptin deficiency confers resistance to diet-induced obesity in mice. Mol Metab 37, 100995 (2020).

62. Ashwood, P. et al. Brief report: plasma leptin levels are elevated in autism: association with early onset phenotype? J Autism Dev Disord 38, 169–175 (2008).

63. Blardi, P. et al. Variations of plasma leptin and adiponectin levels in autistic patients. Neurosci. Lett. 479, 54–57 (2010).

64. Raghavan, R. et al. Fetal and Infancy Growth Pattern, Cord and Early Childhood Plasma Leptin, and Development of Autism Spectrum Disorder in the Boston Birth Cohort. Autism Res 11, 1416–1431 (2018).

65. Singh, U. P. et al. Leptin antagonist ameliorates chronic colitis in IL-10^−^/^−^ mice. Immunobiology 218, 1439–1451 (2013).

66. Fisch, S. et al. Localized Antileptin Therapy Prevents Aortic Root Dilatation and Preserves Left Ventricular Systolic Function in a Murine Model of Marfan Syndrome. J Am Heart Assoc 9, e014761 (2020).

67. Chapnik, N. et al. A superactive leptin antagonist alters metabolism and locomotion in high-leptin mice. J Endocrinol 217, 283–290 (2013).

68. Gonzalez, A. et al. A Leptin Receptor Antagonist Attenuates Adipose Tissue Browning and Muscle Wasting in Infantile Nephropathic Cystinosis-Associated Cachexia. Cells 10, 1954 (2021).

69. Cosentino, L., Vigli, D., Franchi, F., Laviola, G. & De Filippis, B. Rett syndrome before regression: A time window of overlooked opportunities for diagnosis and intervention. Neurosci Biobehav Rev 107, 115–135 (2019).

70. Guy, J., Hendrich, B., Holmes, M., Martin, J. E. & Bird, A. A mouse Mecp2-null mutation causes neurological symptoms that mimic Rett syndrome. Nat Genet 27, 322–326 (2001).

