## supplemental figure for "Leptin antagonism improves Rett syndrome phenotype in symptomatic *Mecp2-*deficient mice"

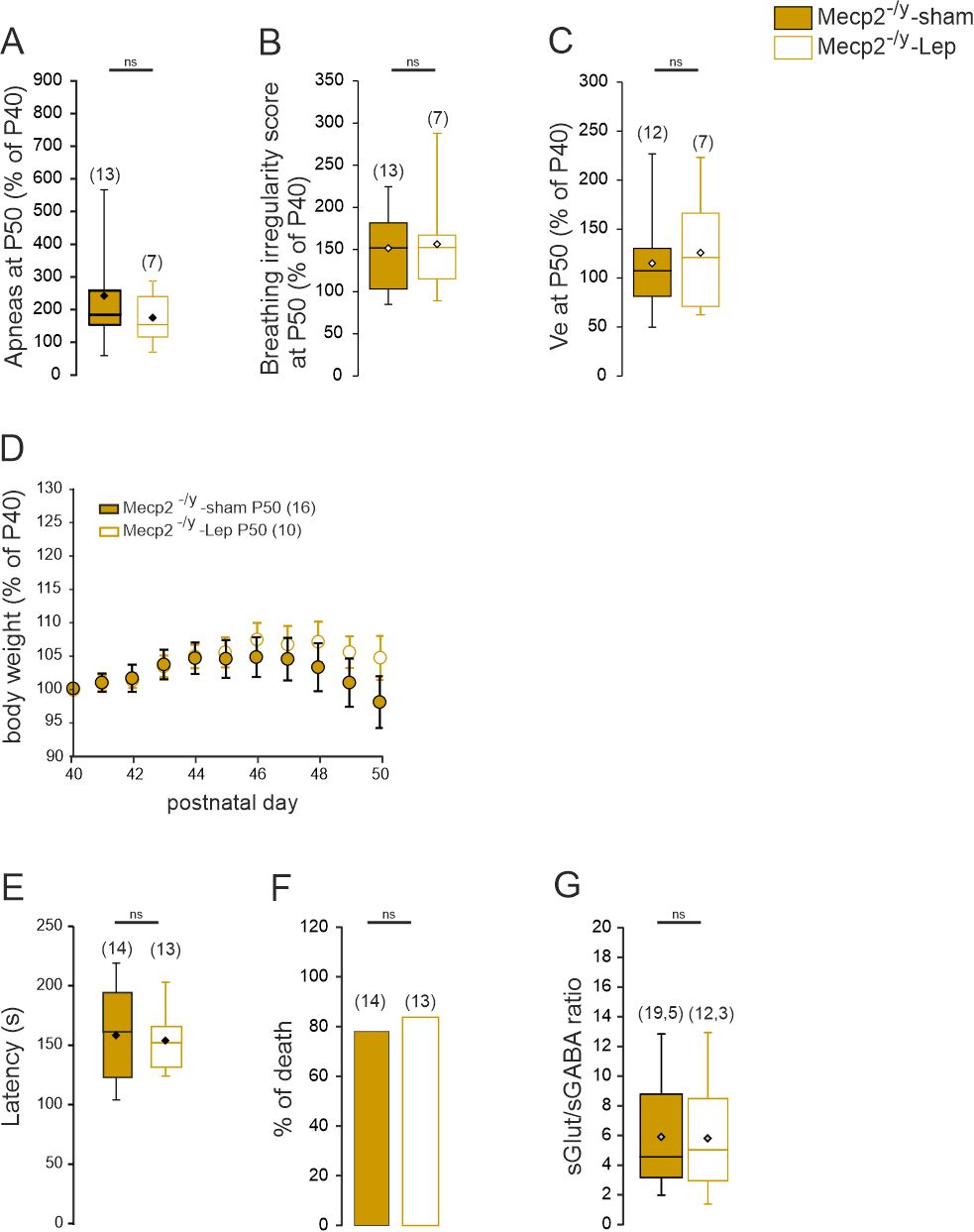

**Supplementary Fig S1: Leptin treatment has no effect on *Mecp2^-/y^* mice.**

P40 *Mecp2^-/y^* received daily sub-cutaneous injection of leptin recombinant (5µg/g) during 10 days. Sham mice received the same volume of vehicle. **A-C**) Box plots of the percentage of change (% of P40) of apnea frequency (**A**), breathing irregularity score (**B**) and minute ventilation (**C**) in sham- and treated- *Mecp2*^-/y^ mice. **D**) Body weight change (% of P40) as a function of age in sham- anti-leptin treated *Mecp2^-/y^* mice. **E)** Box plots of the frequency ratio of spontaneous glutamatergic and GABAergic postsynaptic currents recorded on CA3 pyramidal neurons of sham- and leptin treated- *Mecp2*^-/y^ mice at P50. Numbers in parenthesis indicate the number of mice used. Two-tailed unpaired t-test.

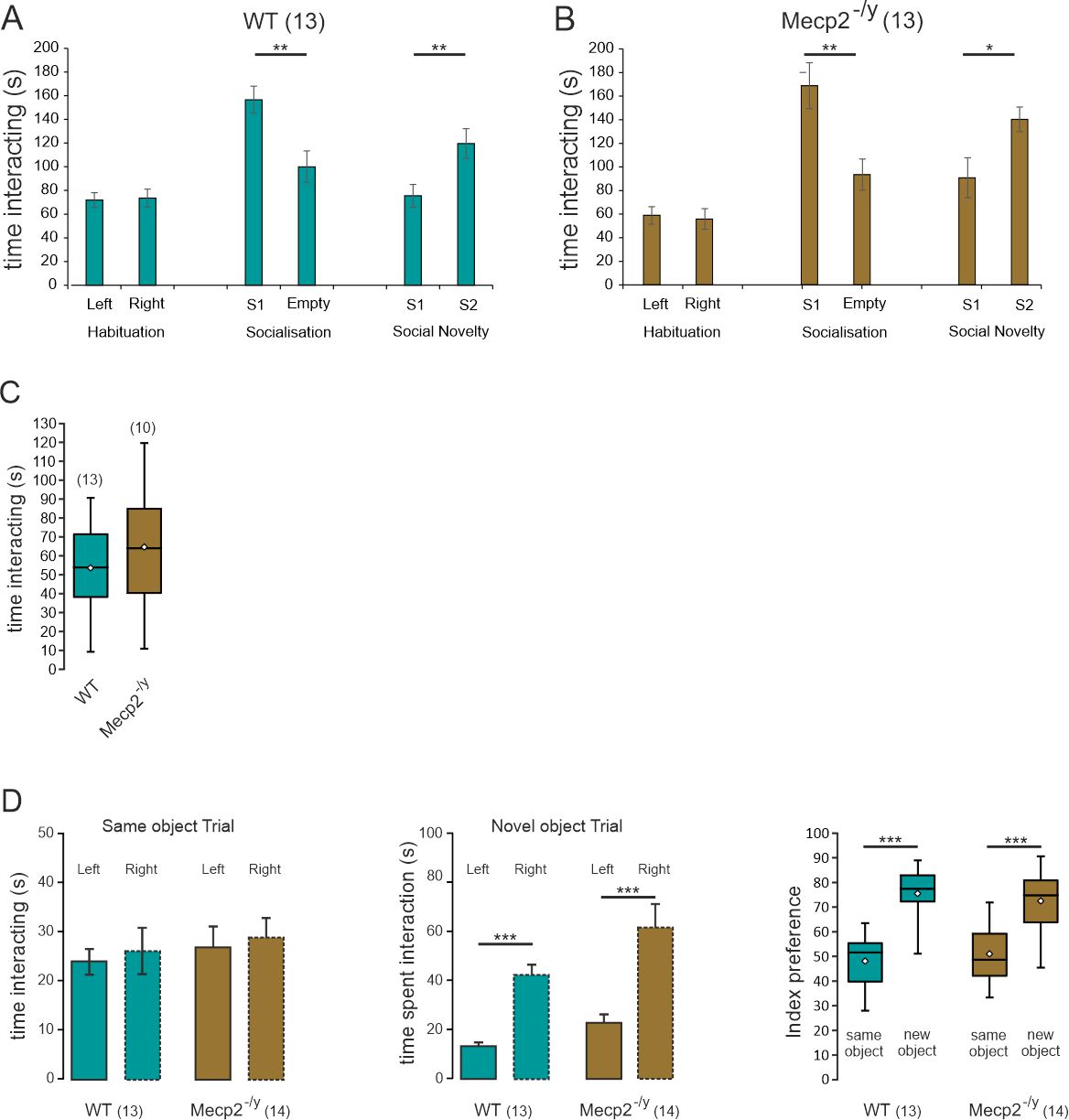

**Supplementary Fig S2: Lack of differences in social and cognitive behavior between P50 WT and *Mecp2^-/y^* mice.**

**A, B**) Mean ± SEM plots of the time interacting of P50 WT (A) and *Mecp2^-/y^* (B) mice during the different phase of the 3 chambers test. **C**) Box plots of the time interacting of P50 WT and *Mecp2^-/y^* mice with a stranger mouse in the spontaneous social interaction test. **D**) Mean ± SEM plots of the time interacting and index preference of P50 WT and *Mecp2^-/y^* mice in the novel object recognition test. Numbers in parenthesis indicate the number of mice used. ***P* < 0.01; ****P* < 0.001, two-tailed paired *t*-test (A, B, D), two-tailed unpaired *t*-test (B, D).

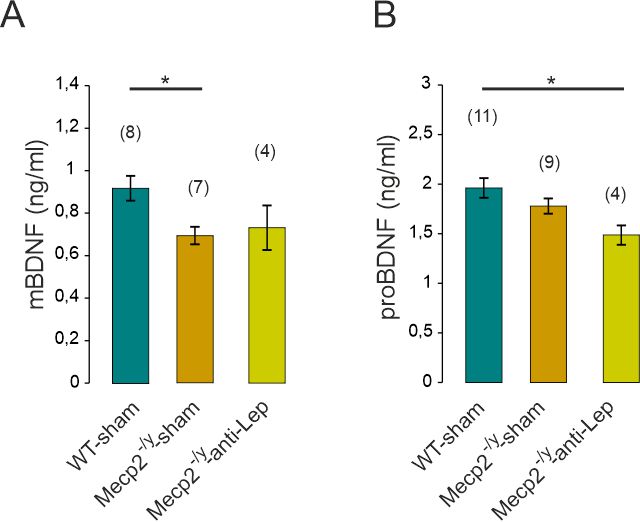

**Supplementary Fig S3: mature and pro-brain derived neurotrophic factor (BDNF) proteins expression in WT and *Mecp2^-/y^* mice.**

**A-C**) Box plots of mature (A) and pro (B) BDNF protein expressions in hippocampal samples taken from P50 WT and *Mecp2^-/y^* mice, using ELISA kit. P40 WT and *Mecp2^-/y^* received sub-cutaneous injection of vehicle (sham) or anti-leptin (5µg/g), every other day, during 10 days. **P* < 0.05, One-way ANOVA followed by a Tukey’s multiple comparison.

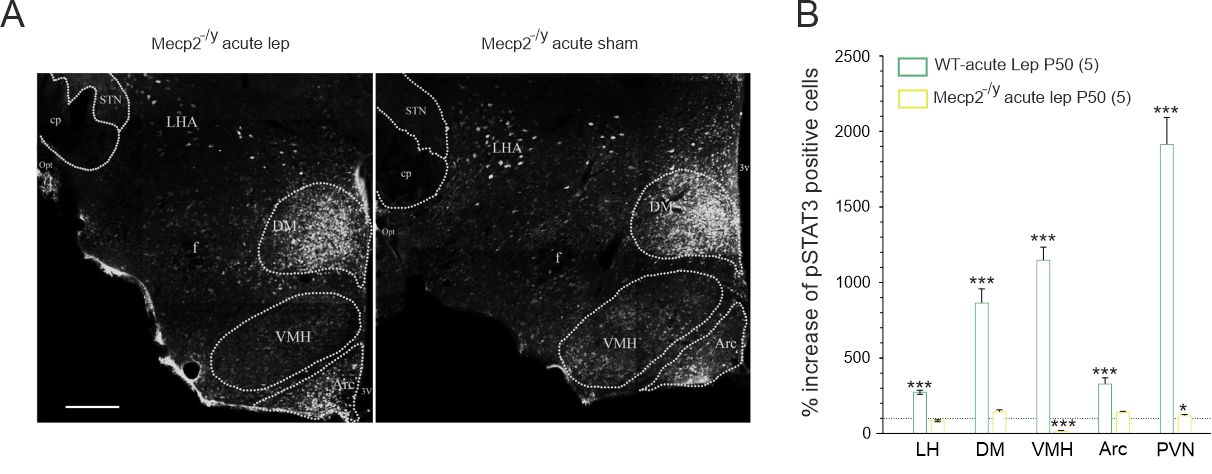

**Supplementary Fig S4: Lack of phenotypic differences between WT and *Mecp2^+/-^* mice.**

**A**) Representative images of pSTAT3 labeling within the arcuate nucleus (Arc) the lateral hypothalamic area (LHA) the ventromedial hypothalamus (VMH) and the dorsomedial hypothalamus (DM) of leptin-treated and sham *Mecp2^-/y^* mice (B) (bregma level -1.94). Note that a high number of pSTAT3 immuno-positive cells, intensely labeled, is detected in the DM and the Arc in both conditions. However, the significant but less intense pSTAT3 labeled cells observed in the VMH of sham *Mecp2^-/y^* mice is no longer observed in leptin-treated *Mecp2^-/y^* mice. 3V: third ventricle; Opt: optic tract; fr: fornix; delineated STN: Subthalamic nucleus; delineated cp: cerebral peduncle, Scale bars, 250 µm. **B**) Percentage of increase of pSTAT3 positive cells induced by sub-cutaneous injection of leptin in WT and *Mecp2*^-/y^ mice. LH: Lateral hypothalamic area; DM: dorsomedial nucleus; VMH: ventromedial nucleus; Arc: arcuate nucleus; PVN: periventricular nucleus.
